# Neuroanatomical abnormalities in a nonhuman primate model of congenital Zika virus infection

**DOI:** 10.1101/2020.11.10.374611

**Authors:** Adele M. H. Seelke, Danielle Beckman, Jeffrey Bennett, Paige Dougherty, Koen K. A. Van Rompay, Rebekah I. Keesler, Patricia A. Pesavento, Lark L. Coffey, John H. Morrison, Eliza Bliss-Moreau

**Affiliations:** California National Primate Research Center, UC Davis, Davis, CA 95616, USA; Department of Psychology, UC Davis, Davis, CA 95616, USA; Department of Neurology, School of Medicine, UC Davis, Davis, CA 95616, USA; Department of Pathology, Microbiology and Immunology, School of Veterinary Medicine, UC Davis, CA 95616, USA

## Abstract

Zika virus (ZIKV) infection of pregnant women can cause major congenital neuronal abnormalities. In the present study, we evaluated neuropathological consequences of fetal ZIKV exposure in rhesus macaques, a highly translatable animal model for human neural development. Quantitative neuroanatomical analyses of the nearly full-term brains of fetuses infected with ZIKV at gestational days 50, 64, and 90, and three procedure-matched sham-inoculated controls were carried out. Whole tissue sections across a complete cerebral hemisphere were evaluated using immunohistochemical and neuroanatomical staining techniques to detect virus localization, identify affected cell types and evaluate gross neuroanatomical abnormalities. None of the subjects were microcephalic. Immunohistochemical staining revealed the presence of ZIKV in the frontal lobe, which contained activated microglia and showed increased apoptosis of immature neurons. ZIKV-infected animals exhibited macrostructural changes within the occipital lobe, including a reduction in gyrification as well as a higher proportion of white matter. Finally, the ZIKV-infected subjects had abnormalities throughout the visual pathway, including disorganization within the lateral geniculate nucleus (LGN) and primary visual cortex (V1). Regional differences tracked with the temporal patterns of the developing brain and likely reflect the neural progenitor cell tropism ZIKV exhibits – painting a picture of inflammatory processes related to viral infiltration sweeping through the cortex, followed by a wave of cell death resulting in morphological changes. These findings may help explain why some infants born with normal sized heads during the ZIKV epidemic manifest developmental challenges as they age, and ultimately may contribute to developing effective treatments and interventions.

**One sentence summary:** Macaque fetuses infected with Zika virus show both macro- and micro-scale neuropathological abnormalities, including decreased gyrencephality, relative increases in cortical white matter, activation of glia, and increased apoptosis.

## Introduction

In late 2015, medical professionals in the northeast region of Brazil reported a surge in the number of children born with microcephaly, with an increase of 265% in the number of cases during 2015-2016 *(1, 2).* Infants born with microcephaly during the Zika outbreak also had a constellation of clinical presentations that ultimately became known as Congenital Zika Syndrome (CZS): skull deformation, abnormally small cerebral cortices, and retinal scarring, among others *(3–5).* Clinicians have identified additional health issues as these children have continued to grow and develop *(6, 7),* including visual impairment *(8),* increased risk for autism *(9),* atypical motor development, increased risk for cerebral palsy *(10),* and poor sleep quality *(11).* Further screening has identified cases where children not diagnosed with CZS at birth exhibit CZS by their first birthday; up to 50% of children born to mothers infected with Zika virus during pregnancy infection exhibit anatomical or behavioral abnormalities *(7, 12).* Taken together, these findings suggest that while microcephaly is an extreme manifestation of CZS, children born with normal sized heads may also have other significant neural deficits. Understanding the anatomy of those deficits is critical for predicting the developmental trajectories of children with CZS and ultimately for developing effective treatments and interventions, and the focus of the present report.

Since the onset of the ZIKV pandemic that began in 2015, a number of studies have revealed important information about how ZIKV targets specific cell types and proteins within the central nervous system (for reviews: *(13, 14)*). The neurotropism of the virus may result from ZIKV using AXL, a tyrosine kinase receptor important in modulating the innate immune system *(15, 16),* to enter the brain *(17–19).* Single-cell RNA sequencing analyses have demonstrated that AXL is highly expressed in radial glia, endothelial cells, microglia and astrocytes in the developing human cortex *(19)*. After breaching the placental-fetal barrier, ZIKV infects radial glial cells. Radial glia cells undergo symmetric and asymmetric cell division to generate the neurons that populate the cerebral cortex *(20, 21)*, and the disruption of this process may ultimately lead to decreased neuronal numbers resulting in microcephaly. Although ZIKV might not be cytotoxic for microglia and astrocytes initially, once infected, these cells can propagate and spread the virus through the brain, maintaining a high viral load in the brain over time *(22)*.

An increasing number of studies point to the involvement of glial cells in ZIKV infection of the CNS. Microglia are well known for their role as cellular effectors of innate immunity; together with astrocytes, they drive the neuroinflammatory process through phagocytosis and cytokine release. They also play a critical role in neural development by promoting neuronal survival via release of neurotrophic factors to support neuronal circuit formation, phagocytosing immature neurons that fail to form proper neuronal circuits, and removing redundant or dysfunctional synapses in the developing brain *(23, 24)*. During the late stages of brain development and after birth, glial cells develop and spread through the nervous system, constituting at least half of the cellular population within the brain *(25)*. Two possible pathways exist by which ZIKV may interact with glial cells –either direct infection or induction of an inflammatory response that activates microglia and/or astrocytes. Congenitally acquired ZIKV has been shown to infect glial precursor cells, impairing their distribution in the brain, leading to reduced and delayed myelination *(26)*, ultimately contributing to neuroinflammation and microcephaly.

Nonhuman primate (NHP) models of ZIKV infection have been established and are able to bridge the translational divide between rodent models and humans *(27, 28).* Rodent models have significant limitations including the fact that adult mouse models of ZIKV must be immunocompromised or genetically modified (e.g. *(29)).* Fetal and neonatal wild type mice do show some susceptibility to ZIKV infection, but they must be directly exposed to the virus, either through intracerebroventricular, intraperitoneal, or intraamniotic inoculation, and as such are not a good model for vertical transmission of ZIKV from infected mothers to fetuses (e.g., *(30)).* In addition, the most severe symptoms of CZS include dysfunction in the neocortex, especially the prefrontal cortex (PFC), and the extent to which certain cortical regions are homologous in rodents and primates is not clear *(31)*. NHPs gestational development is also similar to humans, including its extended duration (relative to rodents), the prevalence of singleton pregnancies (rather than litters), and the structure of the placenta *(32).* The anatomical development and organization of the brain also proceeds along a similar path in humans and NHPs *(33)*. Similar to humans, pregnant NHPs do not need to be immunocompromised to be infected with ZIKV, experience viremia, and demonstrate transplacental transmission of the virus to the fetus *(34–37).* Although none had evidence of microcephaly, previous studies of ZIKV-exposed fetal and infant macaques have identified a number of pathological features, including calcifications, abnormal gliosis and white matter hypoplasia, *(38–41).* Studies that employed immunohistochemical analysis also identified loss of neural precursors, reduced neurogenesis, gliosis and increased apoptosis in different brain regions *(39–41)*. Neuroanatomical evaluations in these studies have adopted approaches that are standard for pathologic analyses, insofar as they typically evaluate small areas of brain tissue and use formalin fixation and paraffin embedding, procedures that can mask epitopes, reduce antigenicity, and make it challenging to understand the impact of infection on the whole brain *(42, 43)*.

The present study builds upon a previously described macaque model of CZS in which, to assure fetal infection at a defined time of gestation, fetuses were inoculated with ZIKV via the intra-amniotic routes, mothers were concurrently inoculated with ZIKV intravenously, and fetuses were harvested at the end of gestation *(40).* While RT-qPCR-based virology, immunology, and histology findings on these animals have been described previously *(40)*, the current report describes a more in-depth neuropathological assessment of the macro- and microstructural effects of ZIKV infection on the developing fetal brain using quantitative microscopy, comparing the effects of ZIKV infection in a developed cortical region (at the caudal extent of the brain, including the occipital lobe) and an immature cortical region (at the rostral extent of the brain, including the frontal lobe). Critically, we maintained the spatial integrity of the tissue and examined whole tissue sections across the complete hemisphere available for study; as such the analyses presented here differ from previous reports because we quantified specific anatomical regions of the brain (e.g., Area 17, Area 46) including features related to their macro-structure (e.g., cortical thickness) and micro-structure (e.g., morphological analyses; identification of immature neurons in the process of apoptosis, etc.). These methods allowed us to identify where the virus was found, what cell types were affected, and whether gross neuroanatomical abnormalities were present.

## Results

To identify and characterize neuroanatomical consequences of fetal ZIKV, we examined the brains of 6 near full-term monkeys, including three animals that had been inoculated with ZIKV at different gestational days (GD 50, 64, and 90; full term is approximately GD 165) and three procedure matched controls *(40).* The pregnant animal inoculated at GD 64 spontaneously gave birth to a small but viable baby on GD 151, at which point both mother and baby were euthanized. The remaining ZIKV-infected subjects and matched controls were monitored until GD 155 when fetectomies were performed followed by necropsy *(40)*. The left cerebral hemisphere was used for both analysis of viral load and histology *(40)* while the right hemisphere was preserved for anatomical analyses. The right hemisphere was sectioned into four blocks to allow for more efficient immersion fixation and cryoprotection (see Figure 1A). Brains were sectioned on a freezing sliding microtome, and one series underwent Nissl staining following standard lab protocols *(44, 45)*. Nissl stained sections from the most anterior block and most posterior block were evaluated to determine if there were gross anatomical abnormalities and if development had proceed as expected (from caudal to rostral; *(33)).* The frontal and occipital lobes were selected as initial targets for evaluation because neocortex develops along the caudal-rostral axis, and at the day of birth the occipital lobe exhibits a well-defined pattern of cortical lamination while the layers in the prefrontal cortex are indistinct *(33, 46).* Additionally, we visually inspected each Nissl stained tissue section to see if there were additional macrostructural abnormalities.

**Figure 1.**
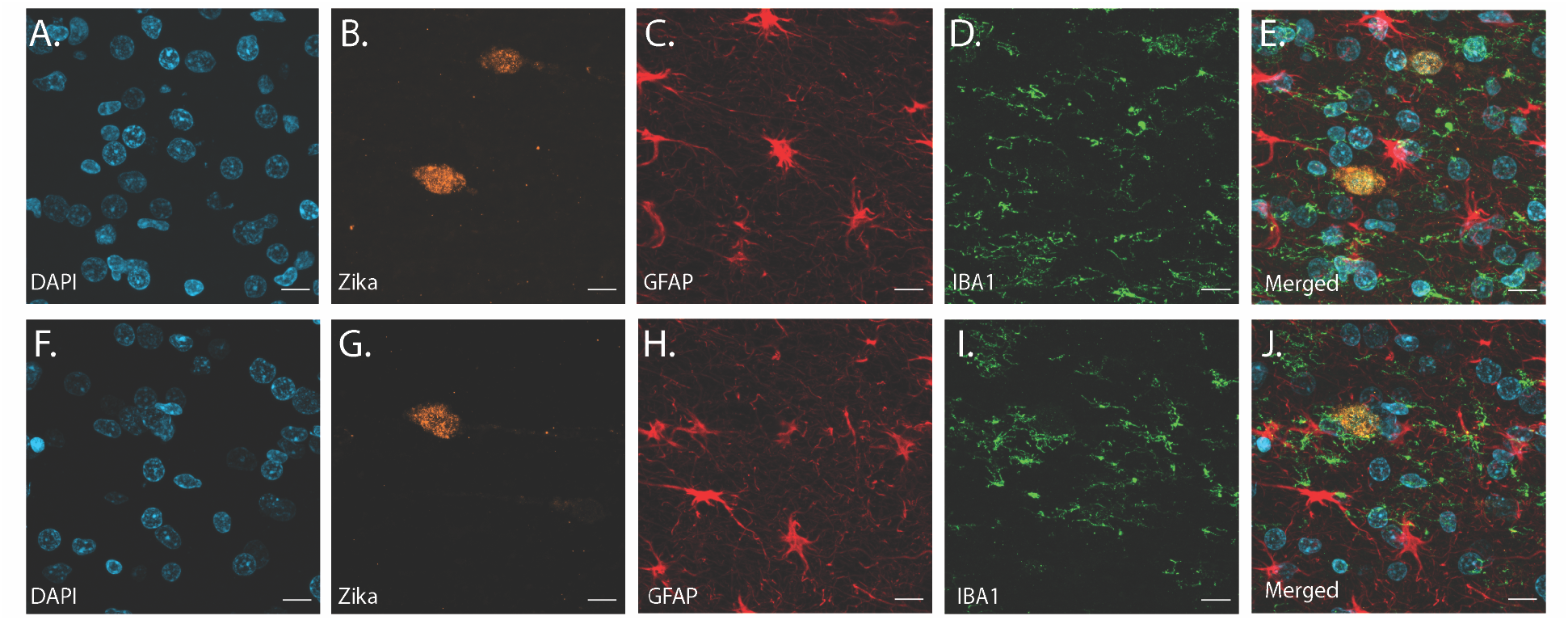
Intraamniotic inoculation of ZIKV results in active brain infections of fetuses up to 105 days post-inoculation. Virus was present in both gray matter (top, A-E) and white matter (bottom, F-J) within the frontal lobe. Sections were stained using immunohistochemistry for DAPI to identify cellular nuclei (A, F), anti-flavivirus group to identify ZIKV envelope (B, G), GFAP to identify astrocytes (C, H), and IBA-1 to identify microglia (D, I). The merged images (E, J) reveal the superimposition of Zika virus envelope over nuclei, along with reactive astrocytes and activated microglia. Scale=10 μm.

### Presence of ZIKV in brains from ZIKV-infected animals

Immunohistochemical (IHC) staining revealed the presence of the ZIKV envelope protein in the ZIKV-infected animals but not in the controls. ZIKV was identified within the gray and white matter of the frontal lobe, surrounded by activated microglia and astrocytes, indicating active infection in those regions (Figure 1). Although viral RNA was sometimes detected by RT-qPCR *(40),* there was no evidence of ZIKV by IHC in any subcortical regions or within the caudal regions of the cortex (data not shown).

### ZIKV Induced Changes to the Occipital and Frontal Lobe

#### Macrostructural Changes

The most prominent manifestation of CZS is microcephaly *(4)*, but none of the ZIKV-infected animals studied here exhibited a disproportionately (compared to their bodies) smaller biparietal diameter during gestation or head sizes at necropsy *(40)*. However, in less severe cases of CZS there have been documented changes to gross anatomical features of the brain including gyral simplification (defined as a reduction in the number of gyri associated with shallow sulci) and lissencephaly. Given those findings in humans, an analysis of the surface-area-to-volume ratio (or gyrencephality) of the occipital and frontal lobes from our infected and control animals allowed for an evaluation of these more subtle (compared to microcephaly) changes related to infection (Figure 2). These analyses revealed important differences in macrostructural features in Zika-infected versus control brains.

**Figure 2.**
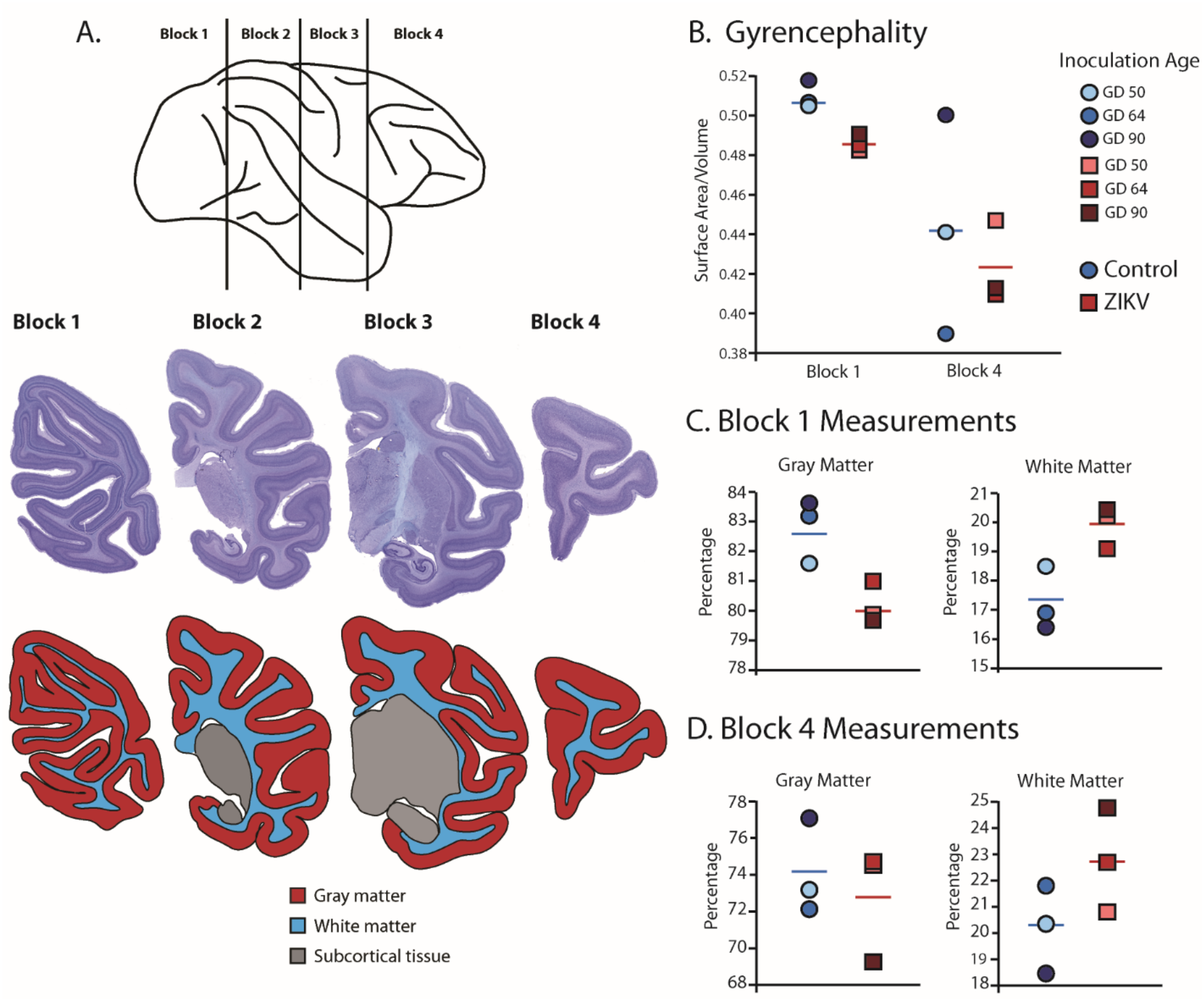
Fetal ZIKV-infection results in variation in cortical development. (A) Each fetal brain was divided into four blocks to ensure complete rapid fixation (top). Representative Nissl stained sections from each block (middle), and the distributions of gray matter (red), white matter (blue), and subcortical tissue (gray) are shown for each section (bottom). For B-D, data from control animals is depicted in blue circles and data from ZIKV-infected animals is depicted in red squares. Gestational day (GD) of infection is indicated with different intensities of blue and red. The mean value for each group is depicted with a horizontal line (blue for control, red for ZIKV). (B) The gyrencephality index of the occipital lobe (Block 1; left) and frontal lobe (Block 4; right) is shown for each individual subject. (C) Proportion of gray matter (left) and white matter (right) in Block 1. (D) Proportion of gray matter (left) and white matter (right) in Block 4.

Gyrification of the occipital lobe was significantly reduced in ZIKV-infected animals as compared with control animals (*t*_4_=3.33, *p*=0.029; *d*=2.72) (Figure 2B). Within the occipital lobes there were no significant differences between ZIKV and control animals in the total lobe volumes (*t*_4_=0.46, p=0.67; *d*=0.37) or white and gray matter volumes (*t*_4_=1.33, *p*=0.25; *d*=1.09 and P=0.264, *p=*0.805; *d*=0.22, respectively). However, differences between the proportions of white and gray matter varied by condition. The occipital lobes from ZIKV animals had a significantly higher proportion of white matter and lower proportion of gray matter than the control animals (*t*_4_=2.93, *p*=0.043; *d*=2.39; because the occipital lobe included only gray and white matter, the proportions are inverses of each other and the statistics identical) (Figure 2C). To evaluate cortical thickness, we selected one clearly defined anatomical region completely contained within the occipital lobe - Brodmann’s Area 17. Cortical thickness in that area did not differ significantly between ZIKV and control subjects (*t*_4_=1.42, *p*=0.230; *d*=1.16; Figure 3A), although all the data points for the ZIKV group were below the mean of the control group, suggesting the possibility of cortical thinning within the primary visual cortex.

**Figure 3.**
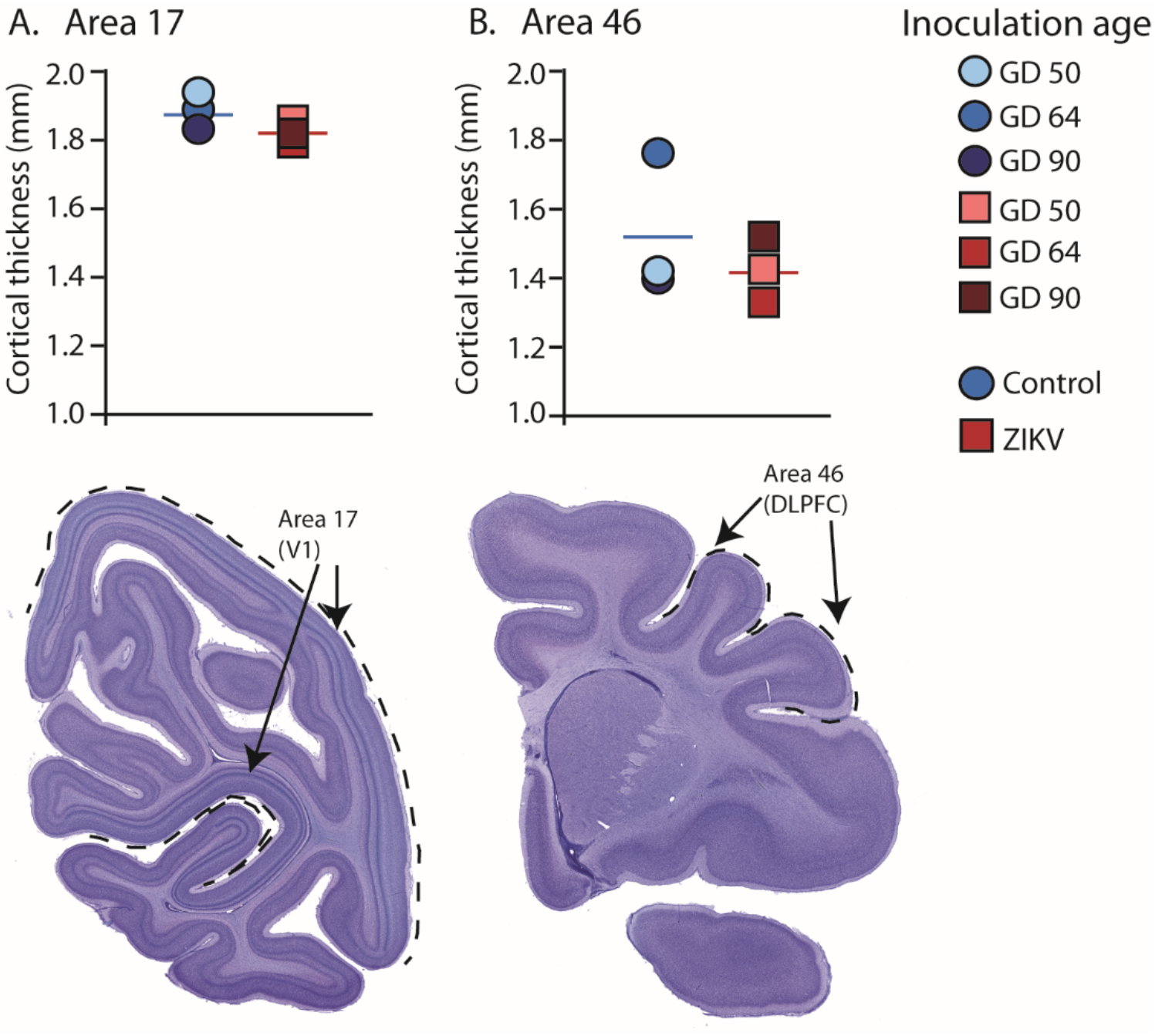
Cortical thickness in Area 17 and 46 did not differ between groups. (A) Cortical thickness of Brodmann’s Area 17 (primary visual cortex, V1) did not differ between the control (blue circles) and ZIKV-infected (red squares) animals. The anatomical localization of Area 17 (primary visual cortex; V1) is shown in a photomicrograph of a section stained for Nissl (dashed lines = the extent of Area 17). (B) Cortical thickness of Brodmann’s Area 46 did not differ between the control (blue circles) and ZIKV-infected animals, although there was more individual variation than was seen in Area 17. The anatomical localization of Area 46 (dorsolateral prefrontal cortex; DLPFC) is shown in a photomicrograph of a section stained for Nissl (dashed line = the extent of Area 46). Conventions as in previous figures.

There was no difference between ZIKV and control animals in the gyrification of the frontal lobes (*t*_4_=0.56, *p*=0.60; *d*=0.46) (Figure 2B). The total volume of the frontal lobes did not differ between ZIKV and control animals *(t4=1.20,p=0.30; d=0.98),* and neither did the white and gray matter volumes (*t*_4_=0.35, *p*=0.75; *d*=0.28 and *t*_4_=1.53, *p*=0.20; *d*=1.25, respectively), nor relative proportions of white and gray matter (*t*_4_=1.80, *p*=0.15; *d*=1.47 and *t*_4_=0.59 *p*=0.59; *d*=0.48, respectively; Figure 2D). As in the occipital lobe analyses, we selected one anatomical region completely contained within the frontal lobe – Brodmann’s Area 46 – and carried out a cortical thickness evaluation of that entire area. There were no significant differences between ZIKV and control animals (*t*_4_=0.57, *p*=0.60; *d*=0.46; Figure 3B). Taken together, these findings suggest that ZIKV infection did not cause gross morphological changes to frontal cortex during development.

#### Microstructural Changes to Glia

Following macrostructural evaluations, we carried out a series of immunohistochemical analyses to quantify cell-level features that might be impacted by ZIKV infection with a specific focus on glia (microglia and astrocytes) in Brodmann’s Area 17 and Area 46.

Changes in glia activation state were measured by analyzing the cell morphology. When glia are activated, their physical shape changes such that its complexity is reduced. Soma increase in size and acquire a rounded shape, a reflection of their transition from ramified ‘resting’ state to an activated ameboid-like phenotype. To analyze glia morphology in details, we performed IHC for IBA-1 (microglial marker) or GFAP (astrocyte marker). Twenty cells from each anatomical area were randomly selected and for each cell a 3D surface rendering was performed to evaluate cell body volume, total cell volume, and number of terminal points.

Analysis of the microglia in Area 17 revealed no significant differences between ZIKV-infected and control animals in terms of cell body volume (*t*_4_=0.74, *p*=0.50; *d*=0.61; Figure 4C) although microglia were marginally smaller in ZIKV-infected animals (whole cell volume) compared to control animals (*t*_4_=2.43, *p*=0.073; *d*=1.97; Figure 4D). There were no group differences in the number of terminal points (*t*_4_=1.78, *p*=0.15; *d*=1.45; Figure 4E). In contrast, microglia in Area 46 of ZIKV-infected compared to control animals, had significantly smaller whole cell volumes (*t*_4_=6.83, *p*=0.0024; *d*=5.58; Figure 4M) and significantly fewer terminal points (*t*_4_=3.41, *p*=0.027; *d*=2.79; Figure 4O). There were, however, no significant group differences in cell body volume (*t*_4_=1.522, *p*=0.203; *d*=1.243; Figure 4N). The smaller whole cell volumes and terminal points suggest that the microglia were in their activated state in frontal cortex. See also Figure S2.

**Figure 4.**
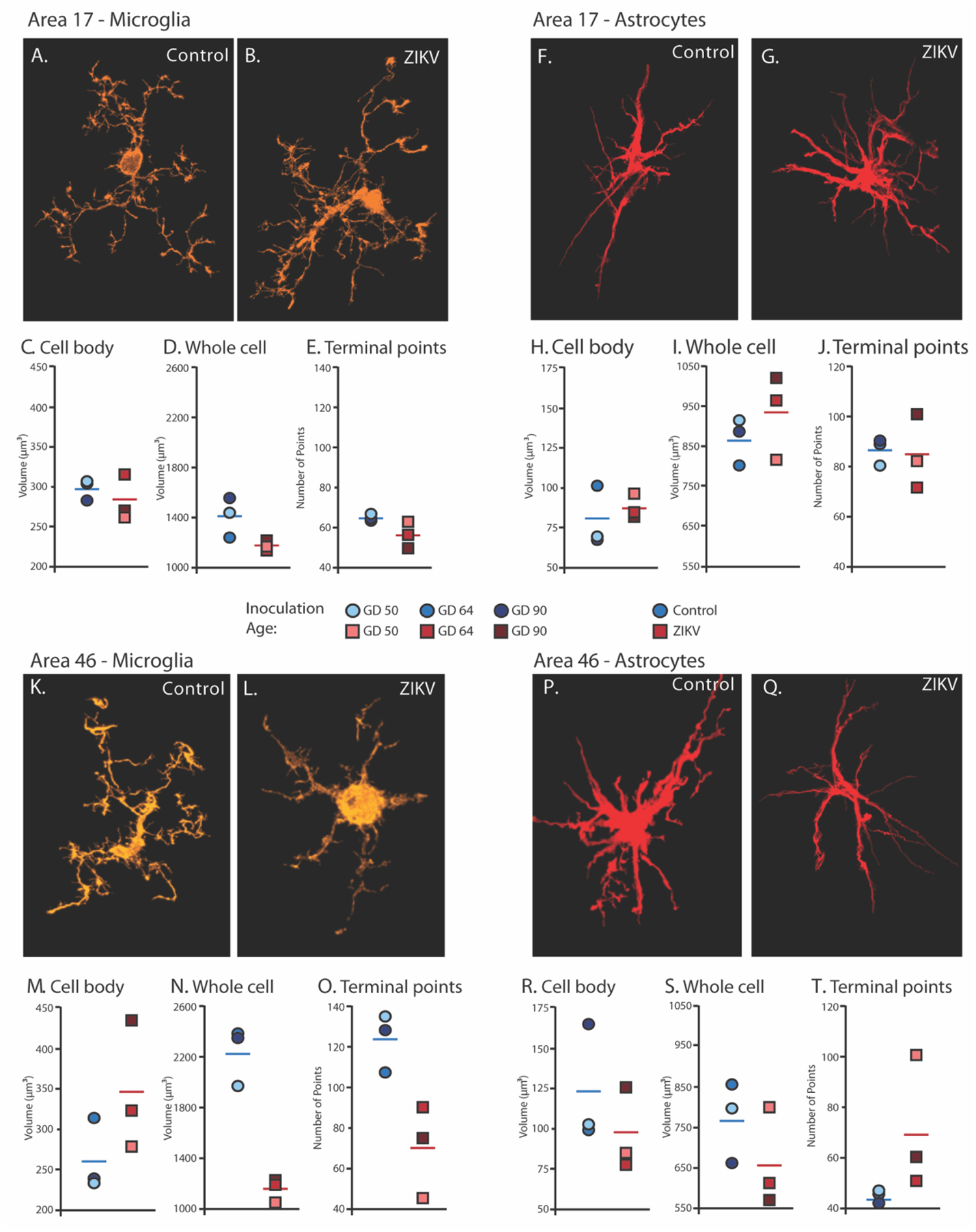
Glial cell morphological changes induced by ZIKV infection of the brain. 3D reconstruction and analysis of glia morphology in Areas 17 and 46 were performed using high-resolution confocal microscopy. Representative microglia in control and ZIKV-infected) animals are depicted in (A) and (B). There were no group differences in cell body volume (C), whole cell volume (D), or number of terminal points (E), a measurement for glia complexity. Similarly, there were not group differences in astrocyte anatomy (F and G for representative anatomy, H-J for data). In contrast, there were notable group differences in Area 46 cellular anatomy (representative images of Area 46 microglia, K and L; astrocytes, P and Q). The cell bodies of ZIKV-infected animals’ microglia were larger in size (M) than control animals. Compared to control animals’ microglia, ZIKV-infected animals’ microglia whole cell volumes were smaller (N) and had fewer of terminal points (O). Compared to controls animals’ astrocytes, ZIKV-infected animals astrocytes tended to have smaller cell body volumes (R), no differences in whole cell volumes (S), and a greater number of terminal points (T).

There were no significant group differences in astrocyte morphology in either Area 17 or Area 46. In Area 17, astrocyte cell body size did not differ between ZIKV-infected and control animals (*t*_4_=0.55, *p*=0.65; *d*=0.43; Figure 4H), whole cell volume (*t*_4_=0.99, *p*=0.38; *d*=0.81; Figure 4I), or number of terminal points (*t*_4_=0.19, *p*=0.86; *d*=0.16; Figure 4J). In Area 46, astrocyte cell body size did not differ between ZIKV-infected and control animals (*t*_4_=1.01, *p*=0.37; *d*=0.83; Figure 4R), whole cell volume (*t*_4_=1.23, *p*=0.29; *d*=1.01; Figure 4S), or number of terminal points (*t*_2.06_= 1.83, *p*=0.21; *d*=1.49; Figure 4T).

#### Microstructural changes related to neural development

The active immune response in the frontal lobe of the brain indicated by the activated microglia (detailed above), in combination with the presence of ZIKV envelope protein, indicated that the frontal lobe was a site of persistent ZIKV infection for a minimum of 60 days post-inoculation. However, the consequences of the infection persist beyond the window of virus replication. Previous studies have demonstrated that ZIKV infection is associated with increased apoptosis of neural progenitor cells (NPCs; *(47, 48)).* IHC fluorescent microscopy analyses for cleaved caspase-3 and SATB2, which are markers of apoptosis onset and immature cortical neurons, respectively, were carried out in order to determine whether NPCs underwent higher rates of apoptosis in immature neurons induced by the presence of ZIKV in the brain in both Area 17 and Area 46. In Area 17 the number of cells in which cleaved caspase-3 and SATB2 were colocalized did not differ across the ZIKV-infected and control groups and the numbers were fairly low (*t*_4_=0.64, *p*=0.55; *d*=0.53; Figure 5A-K), indicating low frequency of immature neuron death. In contrast, there was significantly more frequent death of immature neurons in Area 46 of the ZIKV-infected compared to control animals, as indicated by a greater number of cells that expressed both caspase-3 and SATB2 (*t*_4_=5.37, *p*=0.0057; *d*=4.39; Figure 5L-V; see also Fig S3). The higher frequency of immature neuron death, the active immune response (activated microglia), and the presence of ZIKV protein in Area 46 suggest that frontal cortex remained a site of active ZIKV-induced neuronal remodeling at the time the animals were sacrificed.

**Figure 5.**
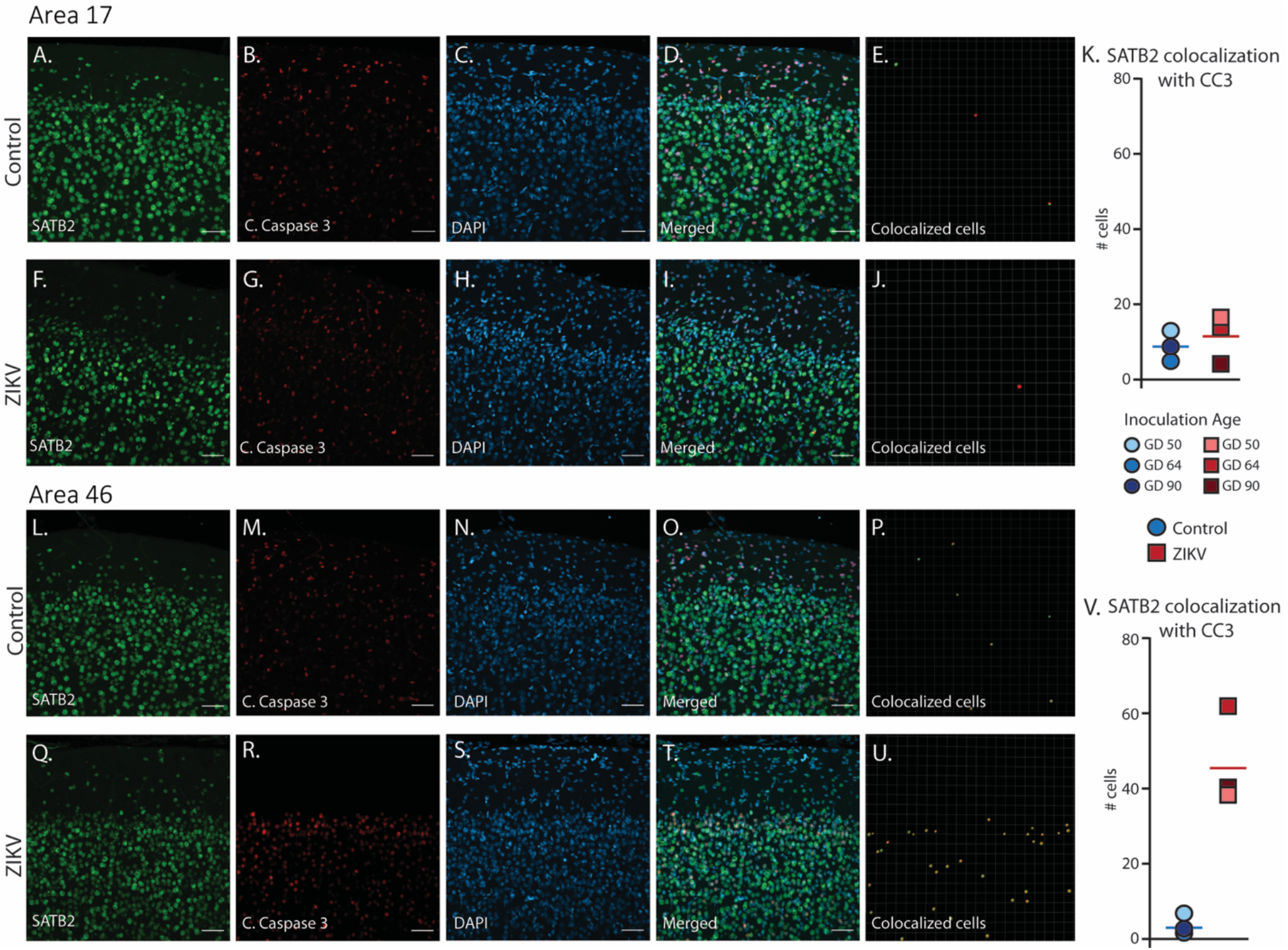
Selective death of immature neurons in Area 46 of ZIKV-infected animals. Representative tissue samples from Area 17 (A-J) and Area 46 (L-U) in ZIKV-infected and control animals. Immature neuronal marker SATB2 (A, F, L, Q) co-localization with cleaved caspase 3 (B, G, M, R), an apoptotic marker, and DAPI (C, H, N, S) were analyzed in Area 17 and 46 of ZIKV-infected animals and CTRs. Merged images (D, I, O, T). 3D surface rendering reconstruction of SATB2 and cleaved caspase 3 are shown in E, J, P, and U. Quantification of co-localization indicated no group differences in the number of immature neurons undergoing apoptosis in Area 17 (K) but a significantly greater immature neurons undergoing apoptosis in ZIKV-infected compared to control animals in Area 46 (V). Scale=25 μm.

#### Additional Neuronal Abnormalities

Visual inspection of the Nissl stained sections of each brain revealed other ZIKV-infection related abnormalities within the visual pathway in addition to those observed in Area 17 (discussed above). These abnormalities included structural changes to the lateral geniculate nucleus (LGN). The LGN is a 6-layered structure that is found in the ventral thalamus that receives retinal input from both eyes via the optic tract *(49)* (for example, Figure 6A). In the ZIKV-infected subjects, we observed multiple instances of laminar discontinuities within the LGN (Figure 6B-D). The LGNs of the control subjects were all normal.

**Figure 6.**
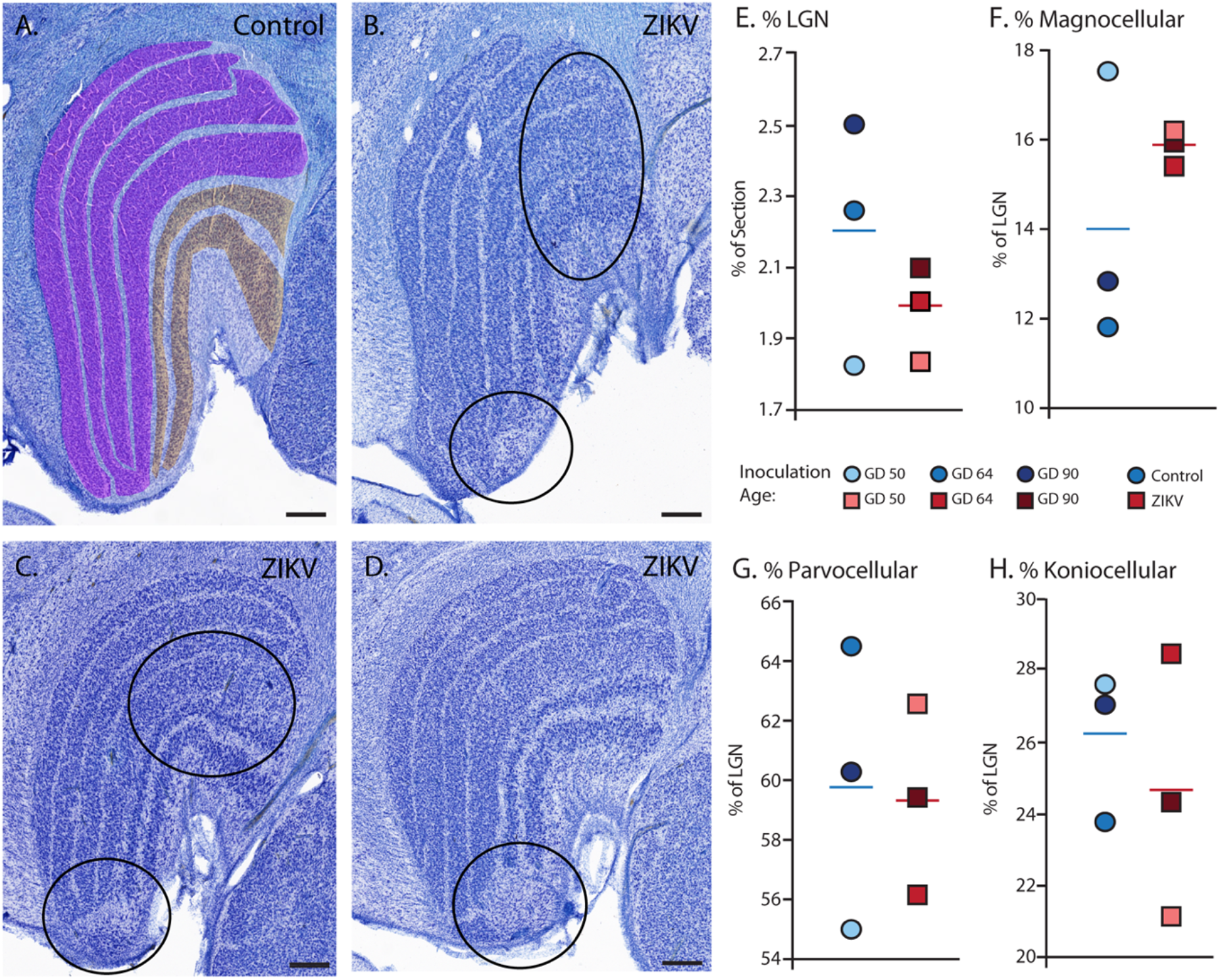
Anatomical abnormalities within the lateral geniculate nucleus (LGN) of ZIKV-infected subjects. (A) An example of normal caudal LGN from the control animal who was sham inoculated at GD90. Layers are highlighted in different colors with the magnocellular layers (1-2) in orange, parvocellular layers (3-6) in magenta; koniocellular layers are in light blue between the other layers. Images (B), (C), and (D) illustrate LGN abnormalities from each of the three subjects infected with ZIKV. Black ovals indicate areas of anatomical abnormalities within each section. (B) Two instances of anatomical abnormalities within the animal infected with ZIKV at GD 50. The top oval highlights an area of undifferentiated lamination, where layers 1, 2, and 3 appear to blend together. The bottom oval highlights an area where the normal laminar structure is interrupted by a region of low cell density. (C) Two instances of anatomical abnormalities within the animal infected with ZIKV at GD 64. The top oval highlights an area of undifferentiated lamination, where layers 2 and 3 appear to blend together. The bottom oval highlights an area where the normal laminar structure is interrupted by a region of low cell density. (D) An instance of laminar interruption of layers 1, 2, and 3 by a region of low cell density in the third ZIKV-infected animal (infected at GD 90). (E) The proportion of the histological section occupied by the LGN did not significantly differ between control (blue circles) animals and the ZIKV-infected animals, *t_4_*=1.00, *p*=0.37, *d*=0.81. There were not group differences in the proportion the (F) Magnocellular (*t*_2.08_=1.02, *p*=0.36, *d*=0.84), (G) Parvocellular (*t_4_*=0.14, *p*=0.90, *d*=0.11), and (H) Koniocellular layers (*t_4_*=0.57, *p*=0.60, *d*=0.47). Scale=250 μm.

The first type of LGN pathology consisted of a blurring of the expected boundaries between cell layers. In healthy tissue, the layers of the LGN are separated by a thin layer of koniocellular tissue (cells between the purple and orange layers in Figure 6A). However, each ZIKV-infected animal had discrete instances where the koniocellular layer was absent, resulting in indistinct boundaries between layers (Figure 6B-D). The location of the restricted absence of the koniocellular layer varied across cases. These abnormalities occurred between both magnocelluar (Figure 6A, purple highlighted area in a control animal) and parvocellular layers (Figure 6A, orange highlighted area in a control animal) and could be found in both the dorsomedial and ventrolateral aspects of the structure.

To investigate the differences observed in the LGN of ZIKV-infected animals, fluorescent microscopy was performed in adjacent sections of the Nissl stained LGN (Figure 7A). In the disrupted LGN layers, the density of the neuronal population (stained with NeuN) in these areas was also substantially reduced (Figure 7B, G, L, and Q) There was an increase in glia activation, mainly in the form of reactive microglia (Figure 7C, G, M, and R), surrounding neurons stained with NeuN (Figure 7B, G, L, and Q) indicating an active inflammatory process in the LGN of the ZIKV-infected animals, although no active virus was observed in this region. No such pathology was observed in the control brains (see Fig S4 in Supplementary Materials).

**Figure 7.**
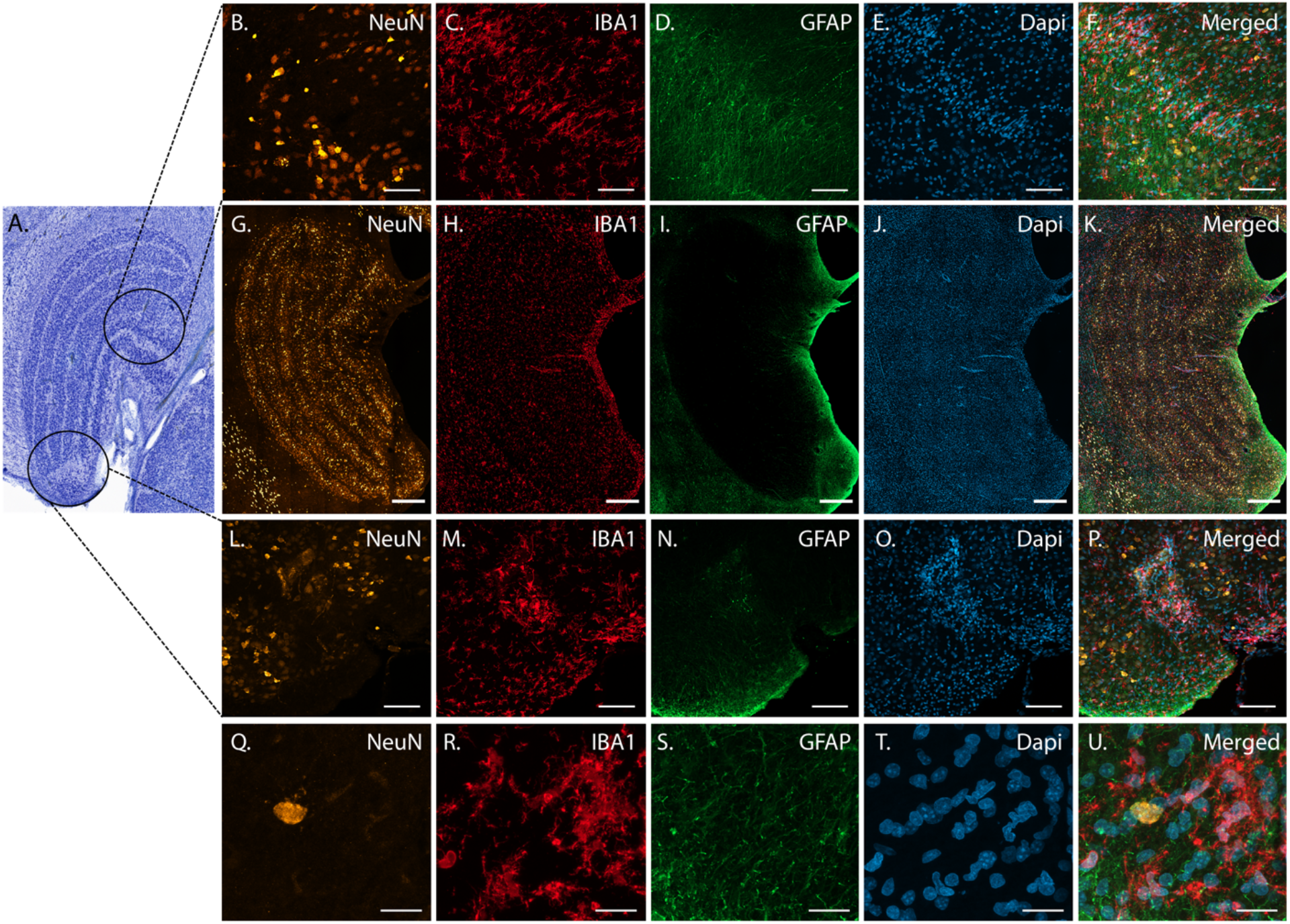
Abnormalities in the LGN of ZIKV-infected animals include reactive glia and neuronal loss. Structural abnormalities were detected in the LGNs of all three ZIKV-infected fetuses in the Nissl stained sections (a representative section is depicted in (A)). A tissue section adjacent to (A) was analyzed using confocal fluorescent microscopy. Neuronal layers were analyzed using neuronal marker NeuN (left column; B, G, L, Q), microglia marker IBA1 (second from left column; C, H, M, R), astrocyte marker GFAP (center column; D, I, N, S) and nuclei marker DAPI (second from right column; E, J, O, T). Merged images are found in F, K, P and U. Scale=250 μm in G-K, scale=25 μm in B-F and L-P, scale=10 μm in Q-U.

The LGN sends direct projections to layer 4 of the primary visual cortex (Area 17) in the occipital lobe. We identified two categories of anatomical abnormalities in the occipital lobe of ZIKV-infected animals (but not controls): cortical heterotopias and laminar disruptions. Cortical heterotopias were identified as ectopic islands of neurons extending into cortical layer 1. Laminar disruptions were identified within Area 17 and consisted of a break in the continuity of layer 4a (Figure 8A-R). These disruptions appeared to be specific to layer 4a and not associated with the normal transition from Area 17 to Area 18, because layers 4b and 4c were still present immediately proximate to the abnormal layer 4a. This disruption is likely functionally significant, as layer 4a receives direct projections from the LGN *(50, 51),* and the absence of cells that make up layer 4a likely represent lost visual information.

**Figure 8.**
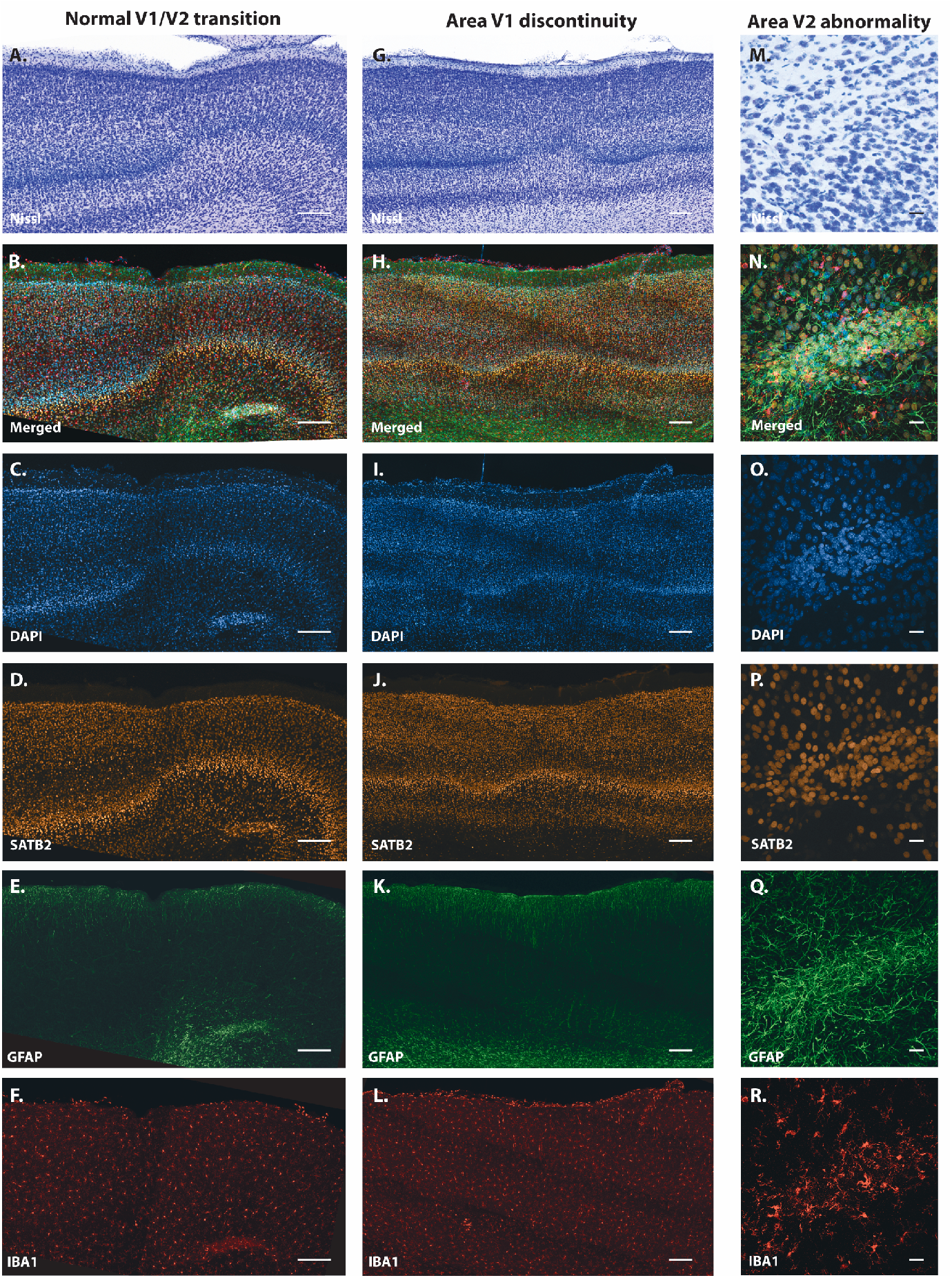
Anatomical abnormalities within the primary and secondary visual cortex of a ZIKV-infected animals. Alterations in primary (V1) and secondary (V2) visual cortex were first detected in the Nissl stained sections (top row; A, G, M). A deeper investigation using quadruple fluorescent staining was performed in adjacent sections for SATB2 (D, J, P), GFAP (E, K, Q) and IBA1 (F, L, R). Merged images are shown in B, H and N. A normal transition between V1 and V2 is characterized by the uniform contraction of layer 4 and the absence of laminar discontinuity (A-F). We identified a region of laminar discontinuity within the boundaries of V1 (G-L) that appears to be the caused by a disruption to layer 4a that resulted in the displacement of layers 4b and 4c towards the pial surface (seen in H, I, J). An additional abnormality was found below layer 6 in area V2 (seen in A-F and magnified in M-R). Disrupted neuronal layers present are surrounded by robust presence of activated microglia and astrocytes, as highlighted in the zoom area N-R. Scale=250 μm in A-L, scale=25 μm in M-R.

## Discussion

Our results demonstrate that ZIKV infection during fetal development results in central nervous system abnormalities that track with temporal patterns of brain development. Importantly, our previous report on these animals *(40)* documented evidence of maternal and fetal virus replication consistent with evidence presented here that ZIKV directly infected the brains of the fetuses. Virus replication and infection-related inflammation were observed in the neocortex of all three ZIKV-infected fetal brains. The types and magnitudes of these abnormalities varied across the cortex, with altered macrostructural features in the occipital cortex and microstructural abnormalities found in the frontal regions of the cortex. These regional differences likely reflect the graded manner in which the neocortex develops, with ZIKV infection tracking temporally with neural development. Importantly, the neuroanatomical abnormalities that we observed in all three ZIKV-infected fetal brains, but not in control brains, occurred even in the absence of the extreme ZIKV infection phenotype documented in human infants, microcephaly. These findings may help explain why some infants born with normal sized heads during the ZIKV epidemic have developed significant sensory, motor, and sociocognitive challenges as they grow up *(6–11).*

The neuroanatomical abnormalities that we identified here appeared to follow the pattern of otherwise normal neuronal development. In mammals the neocortex develops in a gradient. Neural progenitor cells undergo migration in a caudal-to-rostral pattern, resulting in occipital regions having mature laminar organization before mature laminar organization is present in frontal regions *(33, 52).* Rhesus monkeys, like humans and nonhuman primates, have frontal cortices that do not reach maturity until well after birth *(53–55)*. We observed macrostructural differences (e.g., relative grey/white matter volumes) in the occipital lobe which develops first and microstructural and immune-related differences in frontal lobe which develops later. Our analyses revealed evidence that microglia were in their activated state in Area 46 of ZIKV-infected but not control animals, as indicated by smaller whole cell volumes and fewer terminal points. Microglia play a critical role in neural development via a number of mechanisms (including promoting neuronal survival and removing redundant or dysfunctional synapses) and morphological differences across the groups may be evidence of pathological neuronal development. Supporting the idea that ZIKV infection interferes with neural development was our finding that ZIKV-infected fetuses had a greater number of immature cortical neurons undergoing apoptosis compared to control animals. That we found these differences in Area 46, which was still undergoing development at the time of tissue harvest, and not Area 17, which was fully laminated at the time of tissue harvest, underscores the ability of ZIKV to interfere with normative neural development. One interpretation of these data is that at the point at which the tissue was harvested, damage had already been done to the part of the brain that was differentiated but the processes of damage were underway in the tissue that was actively developing. These results suggest that infants infected with ZIKV during fetal development and born with normal sized brains may have neurodevelopmental abnormalities that only become apparent later in development. This finding is of particular importance because most infants born during the 2015-2016 ZIKV epidemic in the ZIKV hotspots were born with normal sized heads and microcephaly, while an extreme phenotype, is thought to be relatively low frequency *(7, 56, 57).*

One seemingly consistent anatomical abnormality found in both human patients and animal models of ZIKV is an increased frequency of abnormalities in the visual system *(58–65),* especially the retina. Patients present with severe eye disease, including congenital cataracts, optic nerve abnormalities, chorioretinal atrophy, intraretinal hemorrhages, mottled retinal pigment epithelium, and blindness *(8, 58, 59, 63–67)*. Further, there is some evidence that ZIKV can be transmitted from the retina to the rest of the visual system via the optic nerve, although this has yet to be confirmed in humans or nonhuman primates *(68)*. Anatomical abnormalities that we observed in the LGN and Area 17/V1 are consistent with infection and developmental disruption of the eye and optic nerve, insofar as projections from the retina enervate the LGN in a retinotopic manner, and projections from the LGN in turn enervate the primary visual cortex (Brodmann’s Area 17) *(69, 70).* The anatomical and functional retinotopic organization of both LGN and V1 is dependent upon activity during developmental critical periods *(71, 72).* Insults during early development that disrupt this activity can result in damage to brain regions downstream of the initial insult *(73–75).* The lesions and laminar dysregulations observed in the LGN and V1, respectively, have significant functional consequences.

While routine H&E histopathology of the eyes did not reveal any significant lesions in these animals *(28)*, subsequent work from our group has identified retinal pathology in one infant that was infected with ZIKV intra-amniotically as a fetus and monitored for 2 years after birth (Yiu et al., submitted). The retinal tissue was used for pathological analysis and was unavailable for use in this study. However, the retinotopic organization and activity-dependent development of LGN layers suggests that the dysregulation we see within the LGN is likely related to lesions of the retina caused by the Zika virus. Future studies would benefit from examination of the entire visual pathway, from retina to LGN to V1, and visual association regions targeted by V1.

While our sample size was small (N=3 in each group), only one hemisphere of the brain was available for our analyses, and the timing of ZIKV infection varied across animals (with one ZIKV and one procedure-matched control at each of three infection dates, GD 50, 64, or 90), all three infected subjects showed the same pattern of pathology: active inflammation in the still-developing regions of the frontal cortex paired with anatomical abnormalities in the already-developed regions of the caudal cortex. Additionally, each infected fetus had evidence of active virus replication as late as 105 days post-inoculation, which could explain the ongoing neurologic damage. Although the sample size here is not atypically small for neuroanatomical analyses of nonhuman primate tissues or initial infectious diseases studies of this sort, it is small and thus important to note that we carried out both experimental and statistical procedures to address potential sample size concerns. First, each infected animal was paired with a procedure-matched control that underwent identical procedures without viral inoculation. Second, all conducted statistical analyses are reported (i.e., there was no p-hacking). Third, raw data but not statistics are displayed in figures encouraging the reader to draw conclusions from the patterns of data within and across groups rather than relying solely on p-values. While replication in a larger sample is a laudable future goal, the present study also sets the stage for targeted hypothesis testing about the timing of infection and gestational development which seems to matter less in the case of ZIKV than potentially anticipated.

Previous studies from our group *(28)* and others *(39, 76–78)* have identified discrete histopathological abnormalities in the brain tissue of macaque fetuses infected with ZIKV, including calcifications with gliosis and a loss of neural precursor cells. In addition to previous neuropathological evaluations, previous evaluation of the other hemisphere of each of these brains identified ZIKV RNA in Area 46 in all infected animals, and in occipital pole (Area 17) of animals infected at GD 64 and GD 90 *(28)*. The histological analyses presented here differ in that we measured specific cytoarchitectonic regions of the brain as well as morphological analyses of glial cells in the frontal and occipital cortex. Together, these analyses paint a picture of inflammatory processes related to viral infiltration sweeping through the cortex, followed by a wave of cell death resulting in morphological changes. Importantly, in addition to underscoring the importance of adopting quantitative neuroanatomical analyses to understand viral infections of the developing brain, these findings underscore the importance of using a monkey model because we would not expect to see such patterns in non-primate models, given that rodents do not have a homolog of Area 46 and a significant period of their brain development occurs after birth.

While we focus heavily on the importance of homologs between humans and rhesus monkeys both in the course of ZIKV infection and the development of prefrontal cortex (including Area 46), the rhesus monkey model opens the possibility of linking neural development to tractable, translatable behavioral development and also developing translatable interventions to improve neural and behavioral function. Rhesus monkeys are a robust model for human infant development across a variety of behavioral domains – including cognition, affect, and social behavior *(79–82).* Moving forward, this will allow us to understand how the neural abnormalities associated with fetal ZIKV infection, such as those documented here, impact infant behavioral development. Because rhesus monkeys develop 3-4 times faster than humans *(33)*, a separate, currently ongoing cohort of infants who were infected with ZIKV as fetuses will soon out-age human infants who were infected *in utero,* thus providing a prospective look at the challenges those infants may face. In this context, our long-term developmental studies which have documented significant neural and behavioral plasticity following early brain damage (e.g., *(83, 84))* provide hope that fairly simple behavioral interventions may be possible to improve the outcomes for impacted children. This is particularly important given that the ZIKV epidemic disproportionally impacted people of color living in socioeconomically disadvantaged communities *(85–87)* and that ZIKV is expected to return seasonally in the future *(88)*.

## Materials and Methods

All procedures were approved by the University of California, Davis Institutional Animal Care and Use Committee which is accredited by the Association for Assessment and Accreditation of Laboratory Animal Care International (AAALAC). Animal care was performed in compliance with the 2011 *Guide for the Care and Use of Laboratory Animals* provided by the Institute for Laboratory Animal Research.

### Study Design

#### Subjects

All adult female rhesus macaques *(Macaca mulatta)* enrolled in the study were from the conventional breeding colony and were born at CNPRC. Animals were determined to be free of West Nile virus (WNV) via simian WNV ELISA (Xpress Bio), SIV-free, type D retrovirus-free and simian lymphocyte tropic virus type 1 antibody negative. Animals were selected from the timed breeding program based on the date of their mating and ultimately the gestational day of their fetus determined by comparing data obtained on ultrasound to CNPRC colony data for fetal development. All animals had at least one and up to six previous successful pregnancies with live births. The animals were housed indoors in standard stainless-steel cages (Lab Products, Inc.) that met or exceed the sizing required by NIH standards, in rooms that maintained 12hr light/dark cycles (0600 lights on to 1800 lights off), with temperature controlled between 65-75°F and 30-70% humidity. Monkeys had free access to water and received commercial chow (high protein diet, Ralston Purina Co.) twice a day and fresh produce multiple times a week. All but one female was paired with a compatible social partner.

#### Experimental Infection with Zika Virus

Four pregnant females and their fetuses were inoculated with a 2015 Brazilian virus isolate (strain Zika virus/H.sapiens-tc/BRA/2015/Brazil_SPH2015; genbank accession number KU321639.1) by injecting 5.0 log10 PFU in 1 ml RPMI-1640 medium both intravenously (IV) and intra-amniotically (IA) *(28).* Each animal was inoculated only once at either 41, 50, 64, or 90 gestational days. The fetus of the female inoculated at GD41 was found dead one week later and its brain was autolyzed *(40).* Three mother-fetus pairs were sham-inoculated with 1ml of RPMI-1640 medium both IV and IA at GD 50, 64, and 90 to serve as procedure-matched controls.

#### Histological Procedures

##### Tissue Harvesting and Fixation

As experimentally planned, the fetuses were harvested at GD 155 via fetectomy and immediately euthanized for detailed tissue collection for 5 of 6 pairs. The fetus that was infected at GD64 was born alive at approximately GD 151 and euthanized the same day for tissue collection. Only evaluation of fetal tissues is reported here.

The left hemisphere of each brain was used for viral load determination or fixed in 10% buffered formalin for histology as described earlier *(40)*. The right hemisphere was fixed in 4% paraformaldehyde in 0.1M Sodium Phosphate buffer for an hour, cut into four blocks to allow sufficient rapid penetration of the fixative into the tissues, and then fixed for 48hrs at 4°C with rotation. Blocks were numbered 1-4, starting at the caudal extent of the brain (Block 1, including the occipital lobe) and ending at the rostral extent of the brain (Block 4, including the frontal pole). The tissue was cryoprotected in 10% glycerin with 2% DMSO in 0.1M sodium phosphate buffer overnight then −20% glycerin with 2% DMSO in 0.1M sodium phosphate buffer for 72hrs *(89).* Blocks were then frozen in isopentane following standard laboratory procedures *(84)*.

##### Tissue sectioning

The tissue was sectioned on a sliding freezing microtome (Thermo Scientific Microm HM430, Waltham, MA USA) into eight series (7 at 30 μm and 1 at 60 μm). 30 μm tissue sections were placed in a cryoprotectant solution and stored at −20°C. The 60 μm tissue sections were postfixed for two weeks in 10% formalin and stored at 4°C.

##### Nissl staining

The 60 μm sections were mounted on gelatin subbed slides and Nissl stained using 0.25% thionin (according to our standard protocols *(45, 84)).* Slides were coverslipped using DPX mounting medium (Millipore Sigma, St. Louis, MO, USA). These sections were scanned (TissueScope LE; Huron Digital Pathology; St. Jacobs, ON, Canada) and digital images were used for analyses. Each image was coded to keep evaluators blind to experimental condition. Anatomical boundaries were determined by comparing Nissl stained tissue to reference atlases *(90, 91)*, and all anatomical analyses were performed using StereoInvestigator (MBF Bioscience; Williston, VT, USA).

##### Immunohistochemistry

30 μm-thick free-floating sections were incubated in an antigen retrieval solution (Wako, S1700) at 60°C for 30 minutes. Sections were incubated in blocking solution: 5% donkey serum, 5% goat serum, 5% BSA in PBS 0.3% triton for 2 hours at RT under agitation. Sections were then incubated overnight with antibodies to: IBA-1 (Wako, code 019-19741, 1:500), NeuN (Synaptic Systems, code: 266-004, 1:1000), GFAP (Abcam, code ab4674, 1:1000), anti-flavivirus group (Millipore, code MAB10216, 1: 400), non-phosphorylated neurofilament (SMI32, Biolegend, code: SMI-32P, 1:500), cleaved caspase 3 (Cell Signaling, code 9661, 1:500) and SATB2 (Cell Signaling, code 39229, 1:500). Tissue was washed thoroughly with PBS and incubated with Alexa Fluor secondary antibodies (Invitrogen, 1:500) for 2 hr, at room temperature. Finally, sections were incubated with DAPI for 10 min after washing the secondary antibodies. Sections were mounted on microscope slides and coverslipped with Prolong Diamond Antifade (Invitrogen).

#### Neuroanatomical Evaluations

We first reviewed each Nissl slide from each case to qualitatively identify potential differences across brains. Given hypotheses about ZIKV’s particular impact on developing neurons, we elected to quantitatively compare Block 1 and Block 4 because of the caudal-rostral developmental gradient of cortical development but also report evaluations of cortical and subcortical structures of interest to our team (e.g., amygdala and hippocampus; see supplemental materials) and those that appeared to be compromised by ZIKV infection (e.g., LGN). Macro-level features of the tissue were quantified on the Nissl sections (including gyrencephalization, cortical thickness, etc., described below). Finally, a series of immunohistochemical analyses on targeted regions of tissue were completed. All people analyzing tissue were blind to condition.

##### Analysis of Nissl stained tissues

To compute an index of gyrencephalization, we measured the area and perimeter of each section of tissue in StereoInvestigator. These areas were chosen because the cortex develops in a caudal to rostral pattern *(52, 92, 93)*; brain development at the time of tissue harvest should have been most complete in the occipital lobe (caudal) and least complete in the frontal lobe (rostral). Thus, if fetal ZIKV infection disrupted development in a timing specific way, would expect to see the greatest difference in neuroanatomical patterning when these lobes were compared. The distance between each tissue section (section thickness + intersection interval) was then used to determine the surface area and volume of each section as follows:

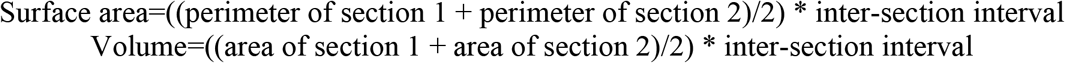

Surface area and volume values were summed for each subject to determine the total surface area and total volume of the frontal and occipital lobes. The gyrencephality ratio for both lobes was computed by dividing the surface area (in mm^2^) by the volume (in mm^3^).

Gray matter and white matter volumes were computed on the same Nissl stained sections in the occipital and frontal lobes. Gray matter was defined as cortical areas that exhibited clear lamination; white matter was defined as fiber tracks adjacent to the cortex. The outline of the section, areas of gray matter, and areas of white matter were traced in StereoInvestigator. Subcortical tissue, including the caudate and putamen and the rostral extent of the hippocampus, was present in a few sections, but these areas were not included in this analysis. The total area and volume of gray and white matter for each section was computed (as above). The proportion of each lobe that was composed of white matter and gray matter was computed by dividing the total volume of gray and white matter in each lobe by the total volume of the lobe. The occipital lobe included no subcortical structures and so the total area was occupied by either gray or white matter. The frontal lobe included subcortical structures and so the percentage of gray matter plus the percentage of white matter did not sum to 100.

Cortical thickness was computed for three areas: Brodmann’s Area 17 (primary visual cortex, V1), Brodmann’s Area 4 (primary motor cortex, M1) (see Supplemental Materials), and Brodmann’s Area 46 (dorsolateral prefrontal cortex). Each region was chosen because of its connection with a sensory input or motor output that is known to be negatively impacted by ZIKV infection and because Area 17 is in Block 1 at the caudal extent of the brain and Area 46 is in Block 4 at the rostral extent of the brain. Humans with CZS have been documented to have vision deficiencies *(8)* as well as muscle spasms and hypertonia *(94)*, implicating V1 and M1, respectively. Similarly, some evidence exists that children with CZS have symptoms consistent with intellectual developmental disorders and executive function disorders that implicate Area 46 *(7, 95)*.

Boundaries of the cortical regions of interest were identified through patterns of cortical lamination and other architectonic landmarks *(90, 91)*. Cortical thickness measurements were taken perpendicular to the pial surface. In Areas 17 and 4, three thickness measurements were taken from each of three adjacent sections. In Area 46, four thickness measurements were taken from each of two adjacent sections. Measurements were first averaged within a section, then between sections, for each subject.

Four sections from each subject that included the full lateral geniculate nucleus (LGN) were evaluated. This procedure was required because the brains were blocked through the LGN in 5 of our 6 subjects. For each subject, the 4 largest complete sections were selected from the available complete sections. Total LGN area was measured on the Nissl stained sections in StereoInvestigator and then the area of magnocellular (layers 1 and 2) and parvocellular (layers 3 through 6) was measured. Koniocellular area for each section was computed as the total area minus the magnocellular and parvocellular areas.

##### Analysis of tissues prepared via immunohistochemical procedures

All analyses of fluorescently stained tissue were conducted using a Zeiss LSM 800 confocal microscope (Carl Zeiss AG, Oberkochen, Germany).

##### Microglia and astrocyte quantification

For microglia and astrocyte morphological analyses, 20 microglia and 20 astrocytes were selected randomly from Area 17 and Area 46 of each animal. For each cell, a z-stack at 63x magnification was made, and the image was exported to Imaris software (Oxford Instruments, Bitplane Inc, Concord, MA, USA) to create 3D volume surface rendering. Cell body volume, whole cell volume, and terminal points were quantified for each cell and averaged within each subject.

##### SATB2 colocalization with cleaved caspase 3

Three z-stack images at 20x magnification were obtained within both Area 46 and Area 17 for each subject. Images that contained both immature neurons (SATB2) and apoptotic cells (cleaved caspase 3) were exported to Imaris software to quantify co-localization.

##### DNA fragmentation

Apoptotic cells were analyzed using DeadEnd™ Fluorometric TUNEL System, from Promega (cat no: G3250). Four slides from each animal were incubated with equilibration buffer for 10 minutes. Slides were then incubated with TUNEL reaction mix for 60 minutes at 37°C in a humidifier chamber, avoiding exposure to light. The reaction consists in a catalytical incorporation of fluorescein-12-dUTP at 3-OH DNA end using Terminal Deoxynucleotidyl Transferase (TdT), forming a polymeric tail using the TUNEL principle. The reaction was stopped using SSC (NaCl and sodium citrate) and slides were washed 3 times with PBS, incubated with DAPI solution for 10 minutes and then mounted using Vectashield (Vector Lab). For each animal, 1 slide was incubated with RQ1 RNase-Free DNase (Prometa cat no M6101) used as a positive control. Three images from each area analyzed were acquired at 20x magnification for each slide for each animal (see Supplemental Materials).

### Data Analysis

Data were analyzed in SPSS 26.0 (IBM Corp). Data were checked for normality via the Kolmogorov-Smirnov and Shapiro-Wilk tests. Data were then analyzed using independent sample t-tests with experimental group as the between subject variable. Cohen’s *d* was computed as a measure of effect sizes. All conducted analyses are reported in the main or supplementary text. We elected to report each individual animals’ data and not to report statistics on the figures because the sample size is small.

## Acknowledgments

Thank you to Jennifer Watanabe, Jodie Usachenko, Christina Cruzen, Kari Christe, Rachel Reader, Wilhelm von Morgenland, Anil Singapuri, Gilda Moadab, and members of the Bliss-Moreau Laboratory who contributed to this work.

## Funding

R21NS104692 and the Murray B. Gardner Junior Faculty Research Fellowship from the UC Davis Center of Comparative Medicine to EBM; R21AI129479 and a pilot grant from the California National Primate Research Center via OD11107 to LLC; and the base grant of the California National Primate Research Center P51OD011107.

## Author contributions

AMHS – carried out histology, wrote manuscript

DB – carried out histology, wrote manuscript

JB – carried out histology, wrote manuscript

PD – carried out histology, edited manuscript

KKAVR – designed and executed study, carried out analysis of viral data, edited manuscript

RIK – designed study, carried out pathology evaluations, edited manuscript

PAP– designed study, carried out pathology evaluations, edited manuscript

LLC – designed and executed study, carried out analysis of viral data, edited manuscript

JHM – oversaw histological analyses, edited manuscript

EBM – designed study, carried out histology, performed statistical analyses, wrote manuscript

## Competing interests

Include any financial interests of the authors that could be perceived as being a conflict of interest. Here also include any awarded or filed patents pertaining to the results presented in the paper.

## Notes

### Competing Interest Statement

The authors have declared no competing interest.

## References

1. Microcephaly Epidemic Research Group, Microcephaly in Infants, Pernambuco State, Brazil, 2015, Emerg. Infect. Dis. 22, 1090–1093 (2016).

2. W. K. De Oliveira, J. Cortez-Escalante, W. T. G. H. De Oliveira, G. M. I. Do Carmo, C. M. P. Henriques, G. E. Coelho, G. V. A. De França, Increase in reported prevalence of microcephaly in infants born to women living in areas with confirmed Zika virus transmission during the first trimester of pregnancy — Brazil, 2015, Morb. Mortal. Wkly. Rep. (2016).

3. C. S. Broussard, C. K. Shapiro-Mendoza, G. Peacock, S. A. Rasmussen, C. T. Mai, E. E. Petersen, R. R. Galang, K. Newsome, M. R. Reynolds, S. M. Gilboa, C. A. Boyle, C. A. Moore, Public Health Approach to Addressing the Needs of Children Affected by Congenital Zika Syndrome HHS Public Access, Pediatrics 141, 146–153 (2018).

4. C. A. Moore, J. E. Staples, W. B. Dobyns, A. Pessoa, C. V. Ventura, E. B. Da Fonseca, E. M. Ribeiro, L. O. Ventura, N. N. Neto, J. F. Arena, S. A. Rasmussen, Characterizing the pattern of anomalies in congenital zika syndrome for pediatric clinicians, JAMA Pediatr. 171, 288–295 (2017).

5. D. Musso, A. I. Ko, D. Baud, D. L. Longo, Ed. Zika Virus Infection — After the Pandemic, N. Engl. J. Med. 381, 1444–1457 (2019).

6. A. C. Wheeler, C. V Ventura, T. Ridenour, D. Toth, L. lia Lima Nobrega, L. Claudia Silva de Souza Dantas, C. Rocha, D. B. Bailey, L. O. Ventura, Skills attained by infants with congenital Zika syndrome: Pilot data from Brazil, PLoS One 13, e0201495 (2018).

7. T. F. Cardoso, R. S. Dos Santos, R. M. Corrêa, J. V. Campos, R. de B. Silva, C. C. Tobias, A. Prata-Barbosa, A. J. L. A. da Cunha, H. C. Ferreira, Congenital Zika infection: neurology can occur without microcephaly., Arch. Dis. Child. 104, 199–200 (2019).

8. L. O. Ventura, C. V. Ventura, L. Lawrence, V. van der Linden, A. van der Linden, A. L. Gois, M. M. Cavalcanti, E. A. Barros, N. C. Dias, A. M. Berrocal, M. T. Miller, Visual impairment in children with congenital Zika syndrome, J. Am. Assoc. Pediatr. Ophthalmol. Strabismus 21, 295–299.e2 (2017).

9. P. Vianna, J. do A. Gomes, J. A. Boquett, L. R. Fraga, J. B. Schuch, F. S. L. Vianna, L. Schuler-Faccini, Zika Virus as a Possible Risk Factor for Autism Spectrum Disorder: Neuroimmunological Aspects, Neuroimmunomodulation 25, 320–327 (2018).

10. F. J. P. Marques, M. C. S. Teixeira, R. R. Barra, F. M. de Lima, B. L. S. Dias, C. Pupe, O. J. M. Nascimento, M. Leyser, Children Born With Congenital Zika Syndrome Display Atypical Gross Motor Development and a Higher Risk for Cerebral Palsy, J. Child Neurol. 34, 81–85 (2019).

11. L. Pinato, E. M. Ribeiro, R. F. P. Leite, T. F. Lopes, A. L. S. Pessoa, L. M. Guissoni Campos, G. E. Piffer, A. L. D. M. Souza, C. M. Giacheti, Sleep findings in Brazilian children with congenital Zika syndrome, Sleep 41, zsy009 (2018).

12. A. C. Wheeler, Development of Infants With Congenital Zika Syndrome: What Do We Know and What Can We Expect?, Pediatrics 141, S154–S160 (2018).

13. S. F. Khaiboullina, F. M. Ribeiro, T. Uppal, E. V Martynova, A. A. Rizvanov, S. C. Verma, Zika Virus Transmission Through Blood Tissue Barriers, Front. Microbiol. 10, 1465 (2019).

14. B. Sutarjono, Can we better understand how zika leads to microcephalyλ A systematic review of the effects of the zika virus on human brain organoids, J. Infect. Dis. (2019), doi:10.1093/infdis/jiy572.

15. C. V. Rothlin, S. Ghosh, E. I. Zuniga, M. B. A. Oldstone, G. Lemke, TAM Receptors Are Pleiotropic Inhibitors of the Innate Immune Response, Cell (2007), doi:10.1016/j.cell.2007.10.034.

16. S. Hafizi, B. Dahlbäck, Signalling and functional diversity within the Axl subfamily of receptor tyrosine kinasesCytokine Growth Factor Rev. (2006), doi:10.1016/j.cytogfr.2006.04.004.

17. L. Meertens, A. Labeau, O. Dejarnac, S. Cipriani, L. Sinigaglia, L. Bonnet-Madin, T. Le Charpentier, M. L. Hafirassou, A. Zamborlini, V. M. Cao-Lormeau, M. Coulpier, D. Missé, N. Jouvenet, R. Tabibiazar, P. Gressens, O. Schwartz, A. Amara, Axl Mediates ZIKA Virus Entry in Human Glial Cells and Modulates Innate Immune Responses, Cell Rep. (2017), doi:10.1016/j.celrep.2016.12.045.

18. F. Rossi, B. Josey, E. C. Sayitoglu, R. Potens, T. Sultu, A. D. Duru, V. Beljanski, Characterization of zika virus infection of human fetal cardiac mesenchymal stromal cells, PLoS One 15, e0239238 (2020).

19. T. J. Nowakowski, A. A. Pollen, E. Di Lullo, C. Sandoval-Espinosa, M. Bershteyn, A. R. Kriegstein, Expression analysis highlights AXL as a candidate zika virus entry receptor in neural stem cells, Cell Stem Cell (2016), doi:10.1016/j.stem.2016.03.012.

20. T. E. Anthony, C. Klein, G. Fishell, N. Heintz, Radial Glia Serve as Neuronal Progenitors in All Regions of the Central Nervous System has shown that they can be divided into multiple antigen-ically distinct subpopulations that differ in their cell cycle kinetics, with each subpopulation changing in a, Neuron 41, 881–890 (2004).

21. S. Falk, M. Götz, Glial control of neurogenesis, Curr. Opin. Neurobiol. 47, 188–195 (2017).

22. J. Muffat, Y. Li, A. Omer, A. Durbin, I. Bosch, G. Bakiasi, E. Richards, A. Meyer, L. Gehrke, R. Jaenisch, Human induced pluripotent stem cell-derived glial cells and neural progenitors display divergent responses to Zika and dengue infections, Proc. Natl. Acad. Sci. U. S. A. (2018), doi:10.1073/pnas.1719266115.

23. M. S. Thion, F. Ginhoux, S. Garel, Microglia and early brain development: An intimate journey Science (80-.). (2018), doi:10.1126/science.aat0474.

24. K. Reemst, S. C. Noctor, P. J. Lucassen, E. M. Hol, The Indispensable Roles of Microglia and Astrocytes during Brain Development, Front. Hum. Neurosci. 10 (2016), doi:10.3389/fnhum.2016.00566.

25. J. B. Zuchero, B. A. Barres, Glia in mammalian development and disease, Dev. (2015), doi:10.1242/dev.129304.

26. L. Chimelli, E. Avvad-Portari, Congenital Zika virus infection: a neuropathological review, Child’s Nerv. Syst. (2018), doi:10.1007/s00381-017-3651-3.

27. S. M. Nguyen, K. M. Antony, D. M. Dudley, S. Kohn, H. A. Simmons, B. Wolfe, M. S. Salamat, L. B. C. Teixeira, G. J. Wiepz, T. H. Thoong, M. T. Aliota, A. M. Weiler, G. L. Barry, K. L. Weisgrau, L. J. Vosler, M. S. Mohns, M. E. Breitbach, L. M. Stewart, M. N. Rasheed, C. M. Newman, M. E. Graham, O. E. Wieben, P. A. Turski, K. M. Johnson, J. Post, J. M. Hayes, N. Schultz-Darken, M. L. Schotzko, J. A. Eudailey, S. R. Permar, E. G. Rakasz, E. L. Mohr, S. Capuano, A. F. Tarantal, J. E. Osorio, S. L. O’Connor, T. C. Friedrich, D. H. O’Connor, T. G. Golos, Highly efficient maternal-fetal Zika virus transmission in pregnant rhesus macaques, PLoSPathog. (2017), doi:10.1371/journal.ppat.1006378.

28. L. L. Coffey, R. I. Keesler, P. A. Pesavento, K. Woolard, A. Singapuri, J. Watanabe, C. Cruzen, K. L. Christe, J. Usachenko, J. Yee, V. A. Heng, E. Bliss-Moreau, J. R. Reader, W. Von Morgenland, A. M. Gibbons, K. Jackson, A. Ardeshir, H. Heimsath, S. Permar, P. Senthamaraikannan, P. Presicce, S. G. Kallapur, J. M. Linnen, K. Gao, R. Orr, T. MacGill, M. McClure, R. McFarland, J. H. Morrison, K. K. A. Van Rompay, Intraamniotic Zika virus inoculation of pregnant rhesus macaques produces fetal neurologic disease, Nat. Commun. 9 (2018), doi:10.1038/s41467-018-04777-6.

29. C. W. Winkler, K. E. Peterson, Using immunocompromised mice to identify mechanisms of Zika virus transmission and pathogenesis Immunology (2018), doi:10.1111/imm.12883.

30. A. M. Paul, D. Acharya, B. Neupane, E. Ashely Thompson, G. Gonzalez-Fernandez, K. M. Copeland, M. Garrett, H. Liu, M. E. Lopez, M. de Cruz, A. Flynt, J. Liao, Y. L. Guo, F. Gonzalez-Fernandez, P. J. S. Vig, F. Bai, Congenital Zika virus infection in immunocompetent mice causes postnatal growth impediment and neurobehavioral deficits, Front. Microbiol. (2018), doi:10.3389/fmicb.2018.02028.

31. M. Laubach, L. M. Amarante, K. Swanson, S. R. White, What, if anything, is rodent prefrontal cortex? eNeuro (2018), doi:10.1523/ENEURO.0315-18.2018.

32. A. M. Carter, Animal Models of Human Placentation - A Review, Placenta (2007), doi:10.1016/j.placenta.2006.11.002.

33. A. D. Workman, C. J. Charvet, B. Clancy, R. B. Darlington, B. L. Finlay, Modeling transformations of neurodevelopmental sequences across mammalian species, J. Neurosci. (2013), doi:10.1523/JNEUROSCI.5746-12.2013.

34. F. Koide, S. Goebel, B. Snyder, K. B. Walters, A. Gast, K. Hagelin, R. Kalkeri, J. Rayner, Development of a zika virus infection model in cynomolgus macaques, Front. Microbiol. (2016), doi:10.3389/fmicb.2016.02028.

35. D. M. Dudley, M. T. Aliota, E. L. Mohr, A. M. Weiler, G. Lehrer-Brey, K. L. Weisgrau, M. S. Mohns, M. E. Breitbach, M. N. Rasheed, C. M. Newman, D. D. Gellerup, L. H. Moncla, J. Post, N. Schultz-Darken, M. L. Schotzko, J. M. Hayes, J. A. Eudailey, M. A. Moody, S. R. Permar, S. L. O’Connor, E. G. Rakasz, H. A. Simmons, S. Capuano, T. G. Golos, J. E. Osorio, T. C. Friedrich, D. H. O’Connor, A rhesus macaque model of Asian-lineage Zika virus infection., Nat. Commun. (2016), doi:10.1038/ncomms12204.

36. T. E. Morrison, M. S. Diamond, Animal Models of Zika Virus Infection, Pathogenesis, and Immunity, J. Virol. (2017), doi:10.1128/jvi.00009-17.

37. L. L. Coffey, P. A. Pesavento, R. I. Keesler, A. Singapuri, J. Watanabe, R. Watanabe, J. Yee, E. Bliss-Moreau, C. Cruzen, K. L. Christe, J. R. Reader, W. Von Morgenland, A. M. Gibbons, A. M. Allen, J. Linnen, K. Gao, E. Delwart, G. Simmons, M. Lanteri, S. Bakkour, M. Busch, J. Morrison, K. K. A. Van Rompay, Zika Virus Tissue and Blood Compartmentaliz-ation in Acute Infection of Rhesus Macaques, PLoS One 12, e0171148 (2017).

38. K. M. Adams Waldorf, J. E. Stencel-Baerenwald, R. P. Kapur, C. Studholme, E. Boldenow, J. Vornhagen, A. Baldessari, M. K. Dighe, J. Thiel, S. Merillat, B. Armistead, J. Tisoncik-Go, R. R. Green, M. A. Davis, E. C. Dewey, M. R. Fairgrieve, J. C. Gatenby, T. Richards, G. A. Garden, M. S. Diamond, S. E. Juul, R. F. Grant, L. Kuller, D. W. W. Shaw, J. Ogle, G. M. Gough, W. Lee, C. English, R. F. Hevner, W. B. Dobyns, M. Gale, L. Rajagopal, Fetal brain lesions after subcutaneous inoculation of Zika virus in a pregnant nonhuman primate, Nat. Med. (2016), doi:10.1038/nm.4193.

39. K. M. Adams Waldorf, B. R. Nelson, J. E. Stencel-Baerenwald, C. Studholme, R. P. Kapur, B. Armistead, C. L. Walker, S. Merillat, J. Vornhagen, J. Tisoncik-Go, A. Baldessari, M. Coleman, M. K. Dighe, D. W. W. Shaw, J. A. Roby, V. Santana-Ufret, E. Boldenow, J. Li, X. Gao, M. A. Davis, J. A. Swanstrom, K. Jensen, D. G. Widman, R. S. Baric, J. T. Medwid, K. A. Hanley, J. Ogle, G. M. Gough, W. Lee, C. English, W. M. I. Durning, J. Thiel, C. Gatenby, E. C. Dewey, M. R. Fairgrieve, R. D. Hodge, R. F. Grant, L. Kuller, W. B. Dobyns, R. F. Hevner, M. Gale, L. Rajagopal, Congenital Zika virus infection as a silent pathology with loss of neurogenic output in the fetal brain, Nat. Med. (2018), doi:10.1038/nm.4485.

40. L. L. Coffey, R. I. Keesler, P. A. Pesavento, K. Woolard, A. Singapuri, J. Watanabe, C. Cruzen, K. L. Christe, J. Usachenko, J. Yee, V. A. Heng, E. Bliss-Moreau, J. R. Reader, W. Von Morgenland, A. M. Gibbons, K. Jackson, A. Ardeshir, H. Heimsath, S. Permar, P. Senthamaraikannan, P. Presicce, S. G. Kallapur, J. M. Linnen, K. Gao, R. Orr, T. MacGill, M. McClure, R. McFarland, J. H. Morrison, K. K. A. Van Rompay, Intraamniotic Zika virus inoculation of pregnant rhesus macaques produces fetal neurologic disease, Nat. Commun. (2018), doi:10.1038/s41467-018-04777-6.

41. M. Mavigner, J. Raper, Z. Kovacs-Balint, S. Gumber, J. T. O’Neal, S. K. Bhaumik, X. Zhang, J. Habib, C. Mattingly, C. E. McDonald, V. Avanzato, M. W. Burke, D. M. Magnani, V. K. Bailey, D. I. Watkins, T. H. Vanderford, D. Fair, E. Earl, E. Feczko, M. Styner, S. M. Jean, J. K. Cohen, G. Silvestri, R. Paul Johnson, D. H. O’Connor, J. Wrammert, M. S. Suthar, M. M. Sanchez, M. C. Alvarado, A. Chahroudi, Postnatal Zika virus infection is associated with persistent abnormalities in brain structure, function, and behavior in infant macaques, Sci. Transl. Med. (2018), doi:10.1126/scitranslmed.aao6975.

42. O. Cinar, O. Semiz, A. Can, A microscopic survey on the efficiency of well-known routine chemical fixatives on cryosections, Acta Histochem. 108, 487–496 (2006).

43. E. G. Jones, B. K. Hartman, Recent advances in neuroanatomical methodology., Annu. Rev. Neurosci. 1, 215–296 (1978).

44. G. W. Kreutzberg, 100 years of nissl staining, Trends Neurosci. (1984), doi:10.1016/S0166-2236(84)80213-1.

45. P. Lavenex, P. B. Lavenex, J. L. Bennett, D. G. Amaral, Postmortem changes in the neuroanatomical characteristics of the primate brain: hippocampal formation, J Comp Neurol 512, 27–51 (2009).

46. J. C. Silbereis, S. Pochareddy, Y. Zhu, M. Li, N. Sestan, The Cellular and Molecular Landscapes of the Developing Human Central Nervous System Neuron (2016), doi:10.1016/j.neuron.2015.12.008.

47. C. Li, D. Xu, Q. Ye, S. Hong, Y. Jiang, X. Liu, N. Zhang, L. Shi, C.-F. Qin, Z. Xu, Zika Virus Disrupts Neural Progenitor Development and Leads to Microcephaly in Mice, Cell Stem Cell (2016), doi:10.1016/j.stem.2016.10.017.

48. B. S. F. Souza, G. L. A. Sampaio, C. S. Pereira, G. S. Campos, S. I. Sardi, L. A. R. Freitas, C. P. Figueira, B. D. Paredes, C. K. V Nonaka, C. M. Azevedo, V. P. C. Rocha, A. C. Bandeira, R. Mendez-otero, R. Ribeiro, M. B. P. Soares, Zika virus infection induces mitosis abnormalities and apoptotic cell death of human neural progenitor cells, Nat. Publ. Gr., 1–13 (2016).

49. E. G. Jones, The Thalamus (Plenum Press, 1985; https://books.google.com/books?id=myULCAAAQBAJ&dq=macaque+lateral+geniculate+edward+jones&source=gbs_navlinks_s).

50. A. M. Thomson, Interlaminar Connections in the Neocortex, Cereb. Cortex (2003), doi:10.1093/cercor/13.1.5.

51. K. D. Miller, Understanding Layer 4 of the Cortical Circuit: A Model Based on Cat V1, Cereb. Cortex (2003), doi:10.1093/cercor/13.1.73.

52. C. J. Charvet, B. L. Finlay, Evo-devo and the primate isocortex: The central organizing role of intrinsic gradients of neurogenesis, Brain. Behav. Evol. 84, 81–92 (2014).

53. J. P. Bourgeois, P. S. Goldman-Rakic, P. Rakic, Synaptogenesis in the prefrontal cortex of rhesus monkeys, Cereb. Cortex 4, 78–96 (1994).

54. Z. Petanjek, M. Judaš, G. Šimić, M. R. Rašin, H. B. M. Uylings, P. Rakic, I. Kostović, Extraordinary neoteny of synaptic spines in the human prefrontal cortex, Proc. Natl. Acad. Sci. U. S. A. 108, 13281–13286 (2011).

55. L. Mrzljak, H. B. M. Uylings, G. G. Van Eden, M. Judáš, Chapter 9 Neuronal development in human prefrontal cortex in prenatal and postnatal stages, Prog. Brain Res. 85, 185–222 (1991).

56. M. F. V. V. Aragao, A. C. Holanda, A. M. Brainer-Lima, N. C. L. Petribu, M. Castillo, V. van der Linden, S. C. Serpa, A. G. Tenório, P. T. C. Travassos, M. T. Cordeiro, C. Sarteschi, M. M. Valenca, A. Costello, Nonmicrocephalic Infants with Congenital Zika Syndrome Suspected Only after Neuroimaging Evaluation Compared with Those with Microcephaly at Birth and Postnatally: How Large Is the Zika Virus “Iceberg”?, Am. J. Neuroradiol. 38, 1427–1434 (2017).

57. A. V. Faiçal, J. C. De Oliveira, J. V. V. Oliveira, B. L. De Almeida, I. A. Agra, L. C. J. Alcantara, A. X. Acosta, I. C. De Siqueira, Neurodevelopmental delay in normocephalic children with in utero exposure to Zika virus, BMJ Paediatr. Open 3, 9–11 (2019).

58. T. S. Aleman, C. V. Ventura, M. M. Cavalcanti, L. W. Serrano, A. Traband, A. A. Nti, A. L. Gois, V. Bravo-Filho, T. T. Martins, C. W. Nichols, M. Maia, R. Belfort, Quantitative assessment of microstructural changes of the retina in infants with congenital Zika syndrome, JAMA Ophthalmol. 135, 1069–1076 (2017).

59. M. P. Fernandez, E. Parra Saad, M. Ospina Martinez, S. Corchuelo, M. Mercado Reyes, M. J. Herrera, M. Parra Saavedra, A. Rico, A. M. Fernandez, R. K. Lee, C. V. Ventura, A. M. Berrocal, S. R. Dubovy, Ocular histopathologic features of congenital Zika syndrome, JAMA Ophthalmol. 135, 1163–1169 (2017).

60. J. J. Miner, A. Sene, J. M. Richner, A. M. Smith, A. Santeford, N. Ban, J. Weger-Lucarelli, F. Manzella, C. Rückert, J. Govero, K. K. Noguchi, G. D. Ebel, M. S. Diamond, R. S. Apte, Zika Virus Infection in Mice Causes Panuveitis with Shedding of Virus in Tears, Cell Rep. 16, 3208–3218 (2016).

61. Y. Simonin, N. Erkilic, K. Damodar, M. Clé, C. Desmetz, K. Bolloré, M. Taleb, S. Torriano, J. Barthelemy, G. Dubois, A. D. Lajoix, V. Foulongne, E. Tuaillon, P. Van de Perre, V. Kalatzis, S. Salinas, Zika virus induces strong inflammatory responses and impairs homeostasis and function of the human retinal pigment epithelium, EBioMedicine 39, 315–331 (2019).

62. M. S. Singh, M. C. Marquezan, R. Omiadze, A. K. Reddy, R. Belfort, W. N. May, Inner retinal vasculopathy in Zika virus disease, Am. J. Ophthalmol. Case Reports 10, 6–7 (2018).

63. C. V. Ventura, M. C. Ventura Filho, L. O. Ventura, Ocular Manifestations and Visual Outcome in Children With Congenital Zika Syndrome, Top. Magn. Reson. Imaging 28, 23–27 (2019).

64. C. V. Ventura, L. O. Ventura, V. Bravo-Filho, T. T. Martins, A. M. Berrocal, A. L. Gois, J. R. De Oliveira Dias, L. Araujo, P. Escarião, V. Van Der Linden, R. Belfort, M. Maia, Optical coherence tomography of retinal lesions in infants with congenital zika syndrome, JAMA Ophthalmol. 134, 1420–1427 (2016).

65. I. Verçosa, P. Carneiro, R. Verçosa, R. Girão, E. M. Ribeiro, A. Pessoa, N. G. Almeida, P. Verçosa, M. B. Tartarella, The visual system in infants with microcephaly related to presumed congenital Zika syndrome, J. AAPOS 21, 300–304.e1 (2017).

66. J. J. Miner, B. Cao, J. Govero, A. M. Smith, E. Fernandez, O. H. Cabrera, C. Garber, M. Noll, R. S. Klein, K. K. Noguchi, I. U. Mysorekar, M. S. Diamond, Zika Virus Infection during Pregnancy in Mice Causes Placental Damage and Fetal Demise, Cell (2016), doi:10.1016/j.cell.2016.05.008.

67. L. O. Ventura, C. V. Ventura, N. de C. Dias, I. G. Vilar, A. L. Gois, T. E. Arantes, L. C. Fernandes, M. F. Chiang, M. T. Miller, L. Lawrence, Visual impairment evaluation in 119 children with congenital Zika syndrome, J. AAPOS (2018), doi:10.1016/j.jaapos.2018.01.009.

68. A. N. van den Pol, G. Mao, Y. Yang, S. Ornaghi, J. N. Davis, Zika virus targeting in the developing brain, J Neurosci 37, 2161–2175 (2017).

69. C. J. Shatz, Impulse activity and the patterning of connections during CNS development, Pattern Form. Phys. Biol. Sci. 5, 299–332 (2018).

70. P. Rakic, Prenatal development of the visual system in rhesus monkey, Philos. Trans. R. Soc. London B Biol. Sci. 278, 245–260 (1977).

71. S. Le Vay, T. N. Wiesel, D. H. Hubel, The development of ocular dominance columns in normal and visually deprived monkeys, J. Comp. Neurol. 191, 1–51 (1980).

72. A. Ghosh, C. J. Shatz, Segregation of geniculocortical afferents during the critical period: A role for subplate neurons, J. Neurosci. 14, 3862–3880 (1994).

73. Y. Ito, M. Shimazawa, Y. Inokuchi, H. Fukumitsu, S. Furukawa, M. Araie, H. Hara, Degenerative alterations in the visual pathway after NMDA-induced retinal damage in mice, Brain Res. 1212, 89–101 (2008).

74. U. T. Eysel, U. Wolfhard, The effects of partial retinal lesions on activity and size of cells in the dorsal lateral geniculate nucleus, J. Comp. Neurol. 229, 301–309 (1984).

75. Y. H. Yücel, Q. Zhang, R. N. Weinreb, P. L. Kaufman, N. Gupta, Effects of retinal ganglion cell loss on magno-, parvo-, koniocellular pathways in the lateral geniculate nucleus and visual cortex in glaucoma, Prog. Retin. Eye Res. 22, 465–481 (2003).

76. A. J. Martinot, P. Abbink, O. Afacan, A. K. Prohl, R. Bronson, J. L. Hecht, E. N. Borducchi, R. A. Larocca, R. L. Peterson, W. Rinaldi, M. Ferguson, P. J. Didier, D. Weiss, M. G. Lewis, R. A. De La Barrera, E. Yang, S. K. Warfield, D. H. Barouch, Fetal Neuropathology in Zika Virus-Infected Pregnant Female Rhesus Monkeys, Cell 173, 1111–1122.e10 (2018).

77. E. L. Mohr, L. N. Block, C. M. Newman, L. M. Stewart, M. Koenig, M. Semler, M. E. Breitbach, L. B. C. Teixeira, X. Zeng, A. M. Weiler, G. L. Barry, T. H. Thoong, G. J. Wiepz, D. M. Dudley, H. A. Simmons, A. Mejia, T. K. Morgan, M. S. Salamat, S. Kohn, K. M. Antony, M. T. Aliota, M. S. Mohns, J. M. Hayes, N. Schultz-Darken, M. L. Schotzko, E. Peterson, S. Capuano, J. E. Osorio, S. L. O’Connor, T. C. Friedrich, D. H. O’Connor, T. G. Golos, Ocular and uteroplacental pathology in a macaque pregnancy with congenital Zika virus infection, PLoS One (2018), doi:10.1371/journal.pone.0190617.

78. R. J. Steinbach, N. N. Haese, J. L. Smith, L. M. A. Colgin, R. P. MacAllister, J. M. Greene, C. J. Parkins, J. B. Kempton, E. Porsov, X. Wang, L. M. Renner, T. J. McGill, B. L. Dozier, C. N. Kreklywich, T. F. Andoh, M. R. Grafe, H. L. Pecoraro, T. Hodge, R. M. Friedman, L. A. Houser, T. K. Morgan, P. Stenzel, J. R. Lindner, R. L. Schelonka, J. B. Sacha, V. H. J. Roberts, M. Neuringer, J. V. Brigande, C. D. Kroenke, A. E. Frias, A. D. Lewis, M. A. Kelleher, A. J. Hirsch, D. N. Streblow, A neonatal nonhuman primate model of gestational Zika virus infection with evidence of microencephaly, seizures and cardiomyopathy (2020).

79. K. A. Phillips, K. L. Bales, J. P. Capitanio, A. Conley, P. W. Czoty, B. A. ’t Hart, W. D. Hopkins, S. L. Hu, L. A. Miller, M. A. Nader, P. W. Nathanielsz, J. Rogers, C. A. Shively, M. Lou Voytko, Why primate models matter, Am. J. Primatol. 76, 801–827 (2014).

80. C. J. Machado, in Building Babies: Primate Development in Proximate and Ultimate Perspective, K. B. H. Clancy, K. Hinde, J. N. Rutherford, Eds. (Springer, New York, New York, 2013), pp. 259–279.

81. S. J. Suomi, in Affective development: A psychobiological perspective, N. Fox, R. J. Davidson, Eds. (Lawrence Erlbaum Associates Publishers, Hillsdale, NJ, 1984), pp. 119–159.

82. S. J. Suomi, in Handbook of attachment: Theory, research, and clinical applications, J. Cassidy, P. R. Shaver, Eds. (The Guilford Press, New York, NY, 1999), pp. 181–197.

83. D. S. Grayson, E. Bliss-Moreau, J. Bennett, P. Lavenex, D. G. Amaral, Neural Reorganization Due to Neonatal Amygdala Lesions in the Rhesus Monkey: Changes in Morphology and Network Structure, Cereb Cortex, 1–14 (2017).

84. E. Bliss-Moreau, G. Moadab, A. Santistevan, D. G. Amaral, The effects of neonatal amygdala or hippocampus lesions on adult social behavior, Behav Brain Res 322, 123–137 (2017).

85. M. J. Moreno-Madriñán, M. Turell, W. Reisen, Factors of concern regarding zika and other aedes aegypti-transmitted viruses in the United States, J. Med. Entomol. 54, 251–257 (2018).

86. L. Kapiriri, A. Ross, The Politics of Disease Epidemics: a Comparative Analysis of the SARS, Zika, and Ebola Outbreaks, Glob. Soc. Welf. 7, 33–45 (2020).

87. E. J. Gómez, F. A. Perez, D. Ventura, What explains the lacklustre response to Zika in Brazil? Exploring institutional, economic and health system context, BMJ Glob. Heal. 3, e000862 (2018).

88. B. Tesla, L. R. Demakovsky, E. A. Mordecai, S. J. Ryan, M. H. Bonds, C. N. Ngonghala, M. A. Brindley, C. C. Murdock, Temperature drives Zika virus transmission: Evidence from empirical and mathematical models, Proc. R. Soc. B Biol. Sci. 285 (2018), doi:10.1098/rspb.2018.0795.

89. D. L. Rosene, N. J. Roy, B. J. Davis, A cryoprotection method that facilitates cutting frozen sections of whole monkey brains for histological and histochemical processing without freezing artifact, J. Histochem. Cytochem. 34, 1301–1315 (1986).

90. K. S. Saleem, N. Logothetis, A combined MRI and histology atlas of the rhesus monkey brain in stereotaxic coordinates (Elsevier/AP, 2012).

91. T. Rohlfing, C. D. Kroenke, E. V. Sullivan, M. F. Dubach, D. M. Bowden, K. A. Grant, A. Pfefferbaum, The INIA19 Template and NeuroMaps Atlas for Primate Brain Image Parcellation and Spatial Normalization, Front. Neuroinform. 6, 00027 (2012).

92. P. Rakic, Pre-and post-developmental neurogenesis in primates, Clin. Neurosci. Res. 2, 29–39 (2002).

93. J. B. Colby, J. D. Van Horn, E. R. Sowell, Quantitative in vivo evidence for broad regional gradients in the timing of white matter maturation during adolescence, Neuroimage 54, 25–31 (2011).

94. A. Pessoa, V. van der Linden, M. Yeargin-Allsopp, M. D. C. G. Carvalho, E. M. Ribeiro, K. Van Naarden Braun, M. S. Durkin, D. M. Pastula, J. T. Moore, C. A. Moore, Motor Abnormalities and Epilepsy in Infants and Children With Evidence of Congenital Zika Virus Infection., Pediatrics 141, S167–S179 (2018).

95. V. van der Linden, A. Pessoa, W. Dobyns, A. J. Barkovich, H. van der L. Júnior, E. L. R. Filho, E. M. Ribeiro, M. de C. Leal, P. P. de A. Coimbra, M. de F. V. V. Aragão, I. Verçosa, C. Ventura, R. C. Ramos, D. D. C. S. Cruz, M. T. Cordeiro, V. M. R. Mota, M. Dott, C. Hillard, C. A. Moore, Description of 13 Infants Born During October 2015–January 2016 With Congenital Zika Virus Infection Without Microcephaly at Birth — Brazil, MMWR. Morb. Mortal. Wkly. Rep. (2016), doi:10.15585/mmwr.mm6547e2.

