## Supplementary Materials for "Neuroanatomical abnormalities in a nonhuman primate model of congenital Zika virus infection"

### Additional evaluations of Nissl stained tissue

#### *M1 cortical thickness, and volumes of the amygdala, hippocampus, and optic tract*

We carried out a few additional gross level analyses of one additional cortical target and three subcortical targets. We evaluated the cortical thickness of one additional cortical area, Area 4, primary motor cortex, was included because it is roughly in the middle of the brain (on the rostral-caudal axis) and thus represents a cortex at a different point in development compared to Area 17 and Area 46. Three thickness measurements were taken on each of three different slides and then the value. There was not a difference in cortical thickness between the Zika and control animals in Area 4 ($t_4$=0.80, $p$=0.47; $d$=0.65; $Mean_{ZIKV}$=2427.76 µm, $SD_{ZIKV}$ =95.64 µm; $Mean_{Control}$=2361.72 µm, $SD_{Control}$=106.62 µm).

Given the senior author's lab's focus on the amygdala and hippocampus, we also analyzed volumes of those structures. There were no differences between the ZIKV and control groups in the volumes of the amygdala, hippocampus. Amygdala: $t_4$=0.095, $p$=0.93; $d$=0.078; $Mean_{Zika}$=79.85 mm$^3$, $SD_{Zika}$ =3.88 mm$^3$; $Mean_{Control}$=79.28 mm$^3$, $SD_{Control}$ =9.45 mm$^3$. Hippocampus:

$t_4$=0.58, $p$=0.59; $d$=0.47; $Mean_{Zika}$=135.82 mm$^3$, $SD_{Zika}$ =10.65 mm$^3$; $Mean_{Control}$=143.97 mm$^3$, $SD_{Control}$=21.84 mm$^3$.

Finally, given the impact of ZIKV on the LGN (documented in the main text) and observations about its impact on the retina, we also carried out a volumetric analysis of the optic tract. There were no group differences between the ZIKV and control animals in the volume of their optic tracts: $t_4$=1.061, $p$=0.35; $d$=0.87; $Mean_{Zika}$=31.24 mm$^3$, $SD_{Zika}$ =6.54 mm$^3$; $Mean_{Control}$=25.09 mm$^3$, $SD_{Control}$=7.62 mm$^3$.

***Additional abnormalities in visual cortex***

In addition to the lamination issues that were documented in visual cortex (and described in the main text), there were a number of instances of suprapial cortical heterotopias in ZIKV-infected, but not control animals, visual cortices. See Fig S1.

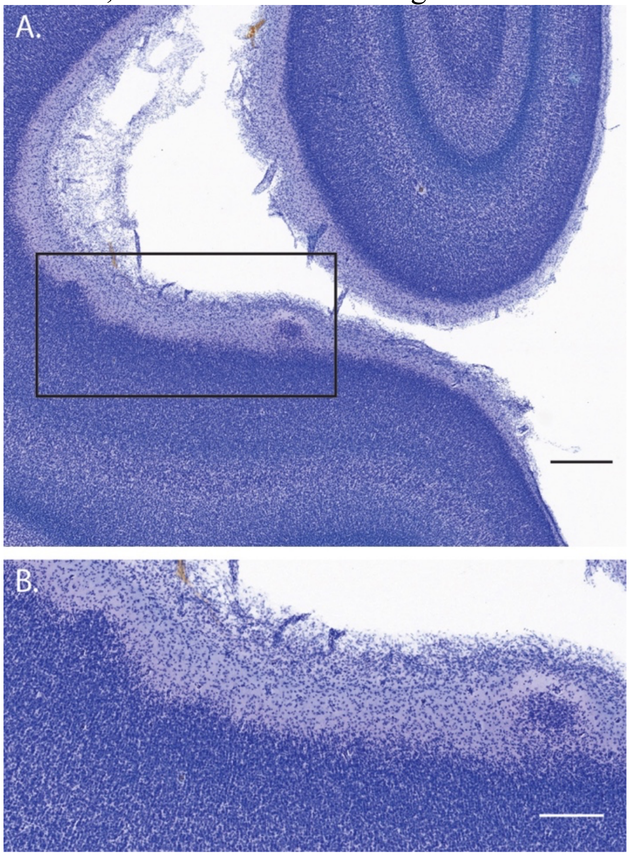


**Fig S1 Suprapial cortical heterotopias observed only in ZIKV-infected animals**. A) A low-magnification photomicrograph of two suprapial cortical heterotopias identified in the cortex of the fetus who was infected with ZIKV at GD90. The black rectangle indicates the region shown in the higher magnification image in B). Scale=500 µm in A, 250 µm in B.

**Additional evaluations of immunofluorescent tissue sections**

We carried out a few additional analyses using multiple labeling fluorescence microscopy to evaluate cell-type specific responses to ZIKV in the brain. Fig S2 depicts common abnormalities found in Area 46 of ZIKV infected animals. Across all three ZIKV-infected animals, clusters of reactive microglia and astrocyte in the deeper cortical layers and disrupted blood vessels were observed (Fig S2A-E). In contrast, non-activated healthy microglia and astrocytes were found surrounding blood vessels and in lower density in Area 46 of control animals (Fig S2F-G).

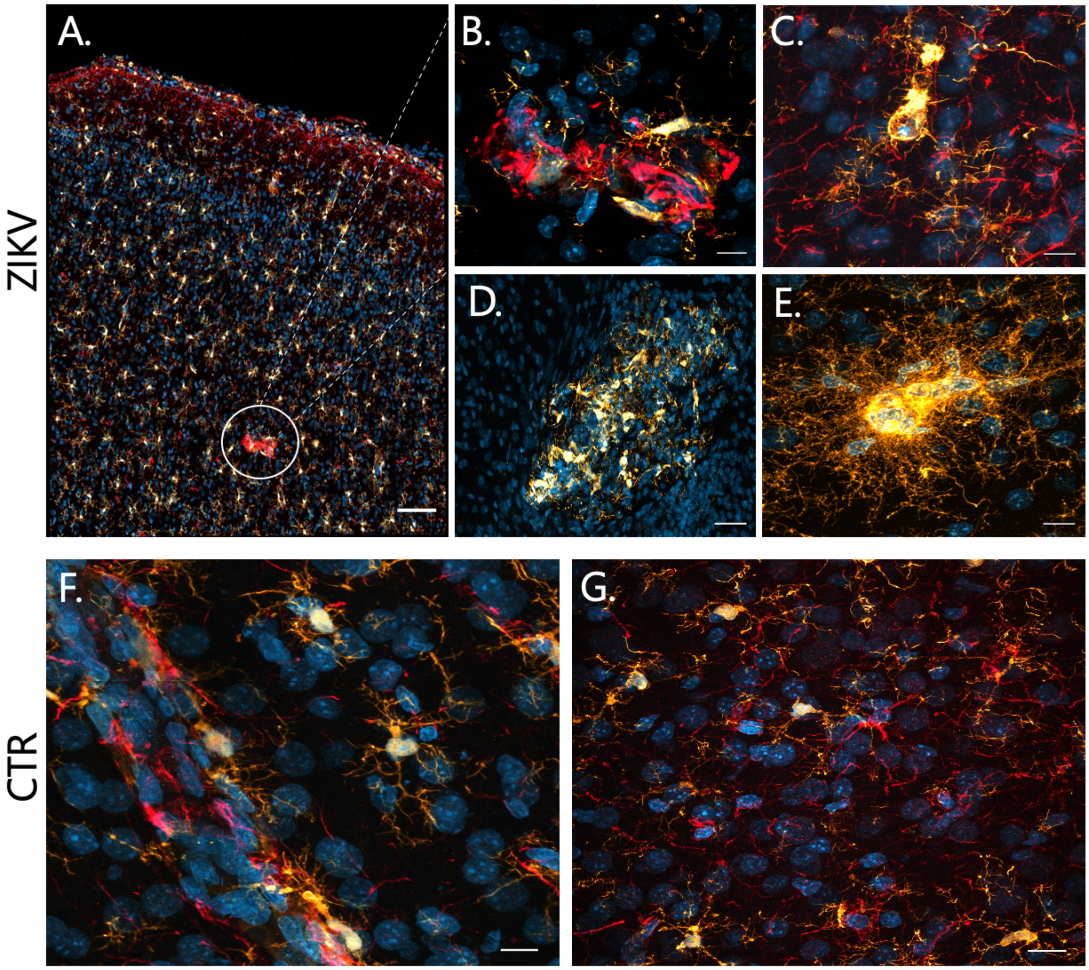


**Fig S2. ZIKV infection induced robust neuroinflammatory response in Area 46.** The animals infected with ZIKV (representative images in A-E) had several cellular abnormalities in Area 46, most consisting of clusters of activated microglia and astrocytes (A-B: DAPI in blue, IBA in yellow, GFAP in red; D-E: DAPI in blue, IBA in yellow). On the other hand, control animals present normal distribution of non-activated microglia and astrocytes surrounding blood vessels and distributed across the cortical layers and white matter (F and G). Scale: 50 μm (A, D), 25 μm (G) and 10 μm (B, C, E, F).

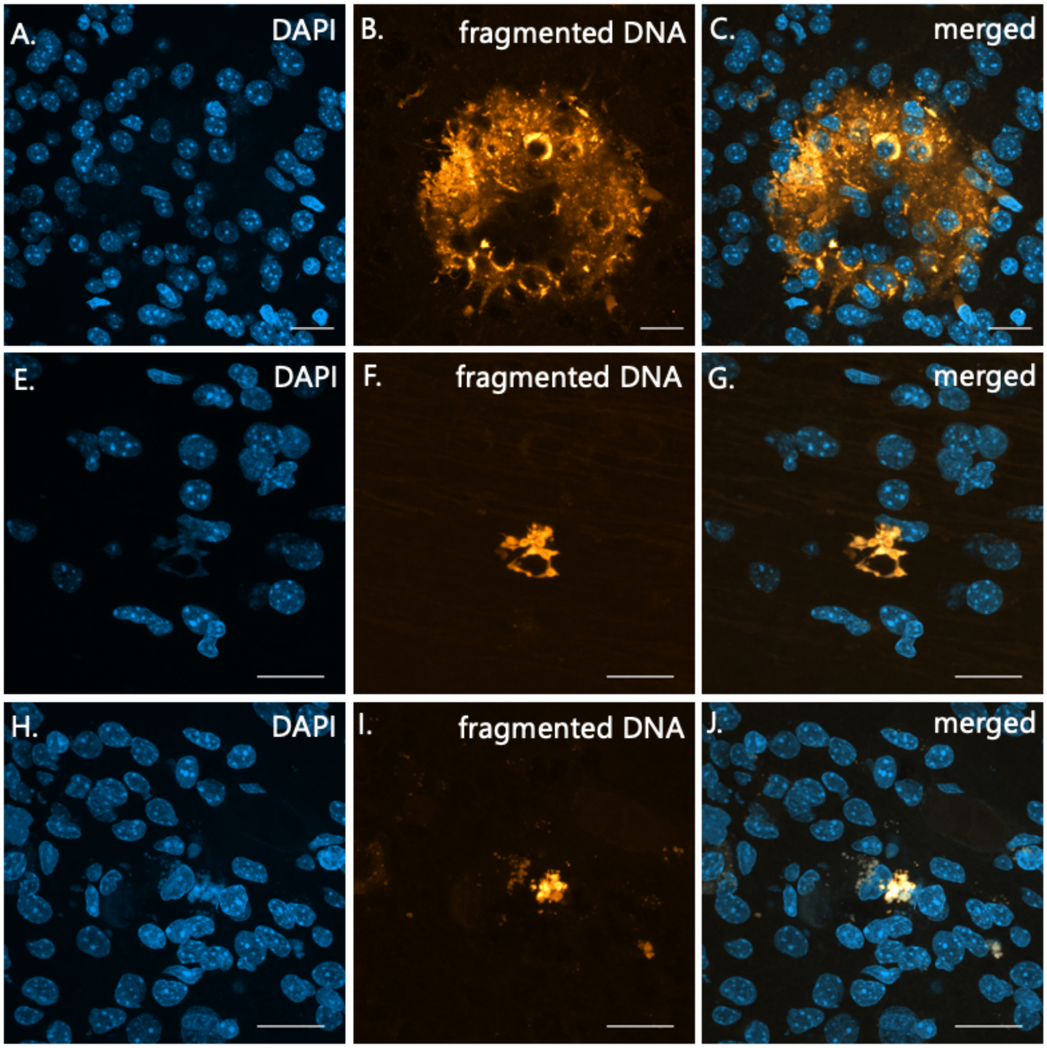


**Fig S3. Abnormal apoptotic clusters in Area 46 of ZIKV-infected animals.** TUNEL assay was performed in tissue slices, with fragmented DNA being detected by fluorescent dye incorporation. Representative images in area 46 (panels A-C and D-F), and in area 17 (panel G-I) highlights the presence of abnormal clusters of fragmented DNA in ZIKV infected animals. Scale: 25 µm (A-C) and 40 µm (D-I).

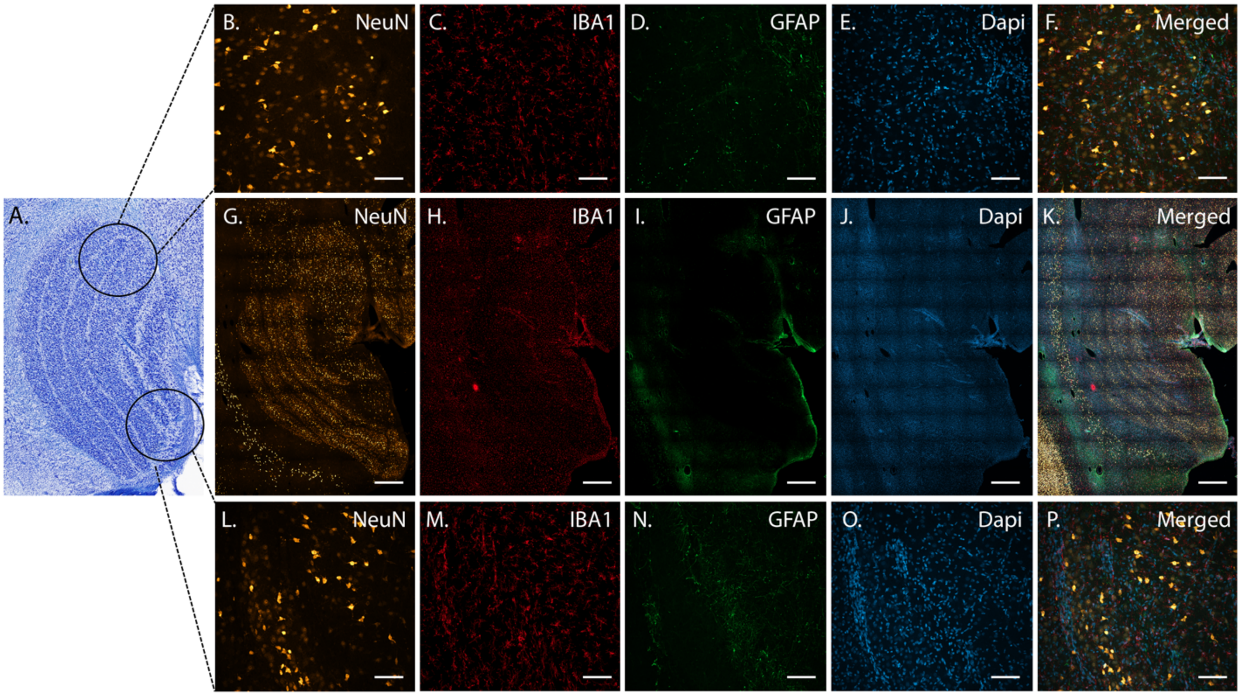


**Supplemental Figure 4. No morphological changes were observed in the LGNs of control animals.** Representative Nissl section of a control animal is shown in A. Analysis of the whole LGN from all control animals were also performed using confocal microscopy. As shown in panel G-K, with randomly picked zoom areas in B-F and L-P, no changes were detected in the pattern of neuronal layers (B, G, L), microglia (C, H, M) and astrocytes (D, I, N) morphology, and DAPI (E, J, O). Merged images are shown in F, K and P. Scale bar: 250 µm (G-K), 25 µm (B-F and L-P).
